# Complex I Governs Iron Levels in Bat Mitochondria to Couple Respiration with Lipid Metabolism

**DOI:** 10.1101/2025.06.10.658522

**Authors:** Weiqian Zeng, Pengwei Peng, Changhan Chen, Dandan Wang, Liuli Cao, Zheng Wang, Yang Xia, Jian Qiu

**Affiliations:** Hunan Key Laboratory of Molecular Precision Medicine, Department of Neurology, Xiangya Hospital, Central South University, Changsha, 410008, China; National Medical Metabolomics International Collaborative Research Center, Central South University, Changsha, 410008, China; National Clinical Research Center for Geriatric Disorders, Xiangya Hospital, Central South University, Changsha, 410008, China; Hunan Key Laboratory of Medical Genetics & Hunan Key Laboratory of Animal Models for Human Diseases, School of Life Sciences, Central South University, Changsha, 410008, China; MOE Key Lab of Rare Pediatric Diseases, Hengyang Medical School, University of South China, Hengyang, 421000, China; Furong Laboratory, Changsha, Hunan, 410008, China

## Abstract

The evolution of mitochondrial respiration has been implicated in the remarkable adaptations of bats, including powered flight, longevity and viral tolerance. However, the mechanisms coordinating mitochondrial respiration with lipid metabolism—a critical energy source for flight—remain poorly understood. Here, we discovered that complex I inhibition triggered iron overload into bat mitochondria, inducing a burst of reactive oxygen species (ROS). This iron translocation relied on the VDAC-MCU axis and mitochondrial membrane potential. Intriguingly, disrupting VDAC oligomerization enhanced iron uptake in a complex I-dependent manner. Mitochondrial iron translocation perturbed fatty acid oxidation and upregulated linoleic acid species, which promoted the reversible accumulation of lipid droplets, suggesting metabolic plasticity crucial for bat energetics. Our findings establish complex I as a pivotal regulatory hub that directly couples respiratory efficiency to lipid metabolism by governing mitochondrial iron levels. This mechanism may represent a key evolutionary adaptation supporting the unique metabolic demands of bat physiology.

**HIGHLIGHTS:**

Complex I inhibition induces unique mitochondrial ROS burst in bat cells.
Mitochondrial iron overload is the primary trigger of complex I-linked ROS burst.
VDAC-MCU axis couples membrane potential to drive mitochondrial iron influx.
Mitochondrial iron translocation remodels lipid metabolism in bat cells.

## INTRODUCTION

Bats represent a unique mammalian lineage that has evolved the remarkable ability to sustain powered flight, a highly energy-intensive mode of locomotion^1^. This evolutionary adaptation has necessitated significant changes in various biological processes, particularly in energy metabolism. The mitochondrial respiratory chain, which is responsible for the production of most cellular ATP through oxidative phosphorylation (OXPHOS), has been a focal point of adaptive evolution in bats^2,3^. Both mitochondrial- and nuclear-encoded genes involved in the OXPHOS system have gone through significant positive selection^4^. These genetic changes in mitochondrial energy metabolism have enabled bats to meet the elevated energy demands required for flight, thus are considered to be crucial for the evolutionary success of bats, allowing them to thrive in diverse ecological niches^3^.

Lipids serve as a dense energy source that can provide the necessary fuel to meet the high energy expenditure associated with flight^5^. The ability to efficiently mobilize and oxidize lipids is essential for the rapid production of ATP to match the high metabolic rate of flight. The catabolism of lipids involves the breakdown of triglycerides into free fatty acids which are subsequently transported into mitochondria to enter the β-oxidation pathway and the tricarboxylic acid (TCA) cycle^6^. This process generates electron donors such as NADH to feed the OXPHOS system for ATP production. However, how mitochondrial respiration is coordinated with lipid metabolism in bat cells remains completely unknown. The intimate regulation of energy production and lipid utilization and storage is crucial for maintaining cellular homeostasis and preventing lipotoxicity^7^.

Iron is an essential element that plays important roles in energy metabolism. It is vital for the biogenesis of iron-sulfur (Fe-S) cluster and heme groups, both of which are integral to the function of the mitochondrial respiratory chain complexes^8^. Mitochondrial complex I (NADH:ubiquinone oxidoreductase), the first and largest enzyme in the electron transport chain, contains multiple Fe-S clusters that are critical for transferring electrons from NADH to ubiquinone^9^. Moreover, iron is a key component of several enzymes involved in the β-oxidation of fatty acids and the TCA cycle^10^. Dysregulation of iron homeostasis can lead to impaired lipid metabolism and mitochondrial dysfunction, resulting in cellular damage and metabolic disorders^11^. Thus, the interplay between iron and lipid metabolism is fundamental to the maintenance of overall cellular health. Whether the adaptive evolution of bat mitochondrial respiration involves concerted regulation of iron homeostasis and lipid metabolism remains enigmatic.

In this study, we perturbed bat respiratory complexes via chemical inhibition and discovered that complex I inhibition unexpectedly triggered large amount of widespread ROS signal (ROS burst), which was stimulated by excessive iron imported into mitochondria. This iron influx depended on mitochondrial membrane potential (ΔΨm) and was regulated by MCU in the inner membrane and VDAC in the outer membrane. The inhibition of VDAC oligomerization further enhanced mitochondrial iron translocation in a complex I-dependent manner. Importantly, mitochondrial iron overload impaired the fatty acid oxidation and caused the buildup of linoleic acid species, promoting the reversible accumulation of lipid droplets. This study provides the first evidence that complex I governs iron levels in bat mitochondria to couple respiration with lipid metabolism.

## RESULTS

### Complex I inhibition induces ROS burst

To perturb mitochondrial respiration in bat cells, various chemicals targeting different complexes of mitochondrial respiratory chain were utilized. Notably, the inhibition of complex I by rotenone induced significant amount of mitochondrial ROS in a concentration dependent manner (Figure 1A). Surprisingly, the highly increased ROS signal was not well confined within mitochondria, but instead spread to other cellular compartments (ROS burst) (Figure 1B). Subsequent quantitative analysis revealed that low concentration of rotenone treatment led to ∼40% of cells with ROS burst, mixed with considerable portion of cells exhibiting increased ROS signal still confined within mitochondria (Figure 1C and 1D). While high concentration of rotenone increased the proportion of cells with ROS burst to more than 80% (Figure 1D). Interestingly, complex I inhibition did not trigger obvious ROS burst in human cells, and only elevated mitochondrial ROS signal slightly (Figure S1A-S1D). Of note, the oxygen consumption rates (basal, ATP-linked and maximal respiratory capacities) of bat cells were significantly lower than those of human cells (Figure S1E and S1F).

**Figure 1.**
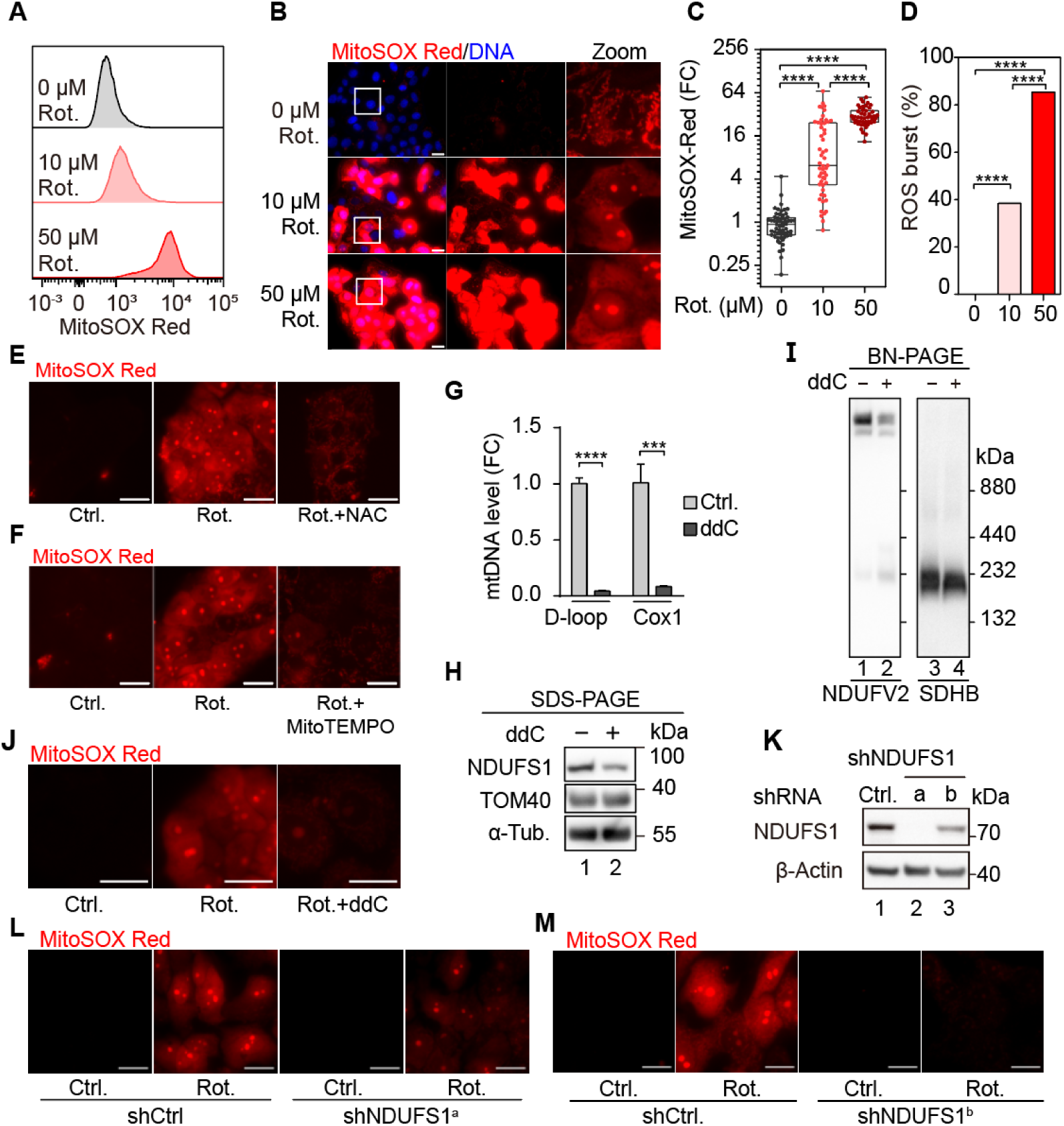
Complex I inhibition induces mitochondrial ROS burst in bat cells. (A) Bat cells (PaKiT) were treated with rotenone (Rot.) at indicated concentration for 6 hours and stained by MitoSOX Red before flow cytometry. (B) Representative images of PaKiT cells treated as in (A). The white squared area was zoomed in with adjusted contrast for clarity. (C and D) Quantification of (B) for signal intensity (C) and percentage of cells with ROS burst (D). FC: fold change. n > 45 and 90 cells for each group in (C) and (D) respectively. (E and F) PaKiT cells were pre-treated with 10 mM N-acetyl cysteine (NAC) (E) or 100 μM MitoTEMPO (F) for 1 day before receiving 50 μM rotenone for 4 hours and imaging for MitoSOX Red. (G) mtDNA of PaKiT was depleted by 100 μM 2′,3′-dideoxycytidine (ddC) for 3 days and analyzed by qPCR (normalized to 18S rRNA, n = 3, mean ± s.d.). (H and I) Protein lysates of PaKiT treated as in (G) were separated by SDS-PAGE (H) or BN-PAGE (I) followed by immunoblotting with indicated antibodies. (J) PaKiT cells were pre-treated with 100 μM ddC for 2 days followed by 50 μM rotenone treatment for 6 hours before imaging of ROS signal. (K) NDUFS1 was knocked down by shRNA for 3 days in PaKiT cells before SDS–PAGE and immunoblotting with indicated antibodies. (L and M) NDUFS1 was knocked down by shRNA for 2-3 days in PaKiT cells before 50 μM rotenone treatment for 1h and imaging of ROS signal. Statistical analysis was performed using one-way ANOVA with Tukey’s test for multiple comparisons for (C), Fisher’s exact test for (D) and two-tailed Student’s *t*-test for (G), \*\*\**P* < 0.001, \*\*\*\**P* < 0.0001. Scale bar = 20 μm. See also Figure S1.

The high level of ROS signal induced by complex I inhibition could be scavenged by N-acetyl cysteine (NAC) and MitoTEMPO (Figure 1E and 1F). The assembly of complex I is coordinated by subunits expressed from both mitochondrial and nuclear genomes^9^. To investigate whether the widespread ROS signal originated truly from bat mitochondria, mitochondrial DNA (mtDNA) was depleted by 2′,3′-dideoxycytidine (ddC) (Figure 1G). As expected, the expression of core components (including NDUFS1) and the assembly of complex I was suppressed by ddC (Figure 1H and 1I). Importantly, ddC treatment effectively inhibited the ROS burst induced by rotenone (Figure 1J), supporting the mitochondrial origin of the high-level ROS. To further investigate whether the ROS burst was complex I dependent, the expression of NDUFS1 was knocked down by two different shRNAs and the ROS burst induced by rotenone in bat cells was effectively inhibited (Figure 1K-1M). Taken together, we conclude that complex I inhibition by rotenone induces mitochondrial ROS burst in bat cells.

### Iron homeostasis is required for ROS burst

To dissect the regulatory mechanisms of ROS burst upon complex I inhibition, we searched for chemicals that could inhibit the ROS signal. Unexpectedly, iron chelators 2,2’-bipyridine (Bpy) and deferoxamine (DFO) both effectively inhibited the ROS burst in bat cells treated with rotenone (Figure 2A, 2B and S2A), indicating the central role of iron homeostasis in the ROS burst event of bat cells. Indeed, supplementing iron in culture medium further promoted the ROS production (Figure S2B). Iron, an essential element for life, is critical for iron-sulfur cluster assembly and heme synthesis in mitochondria^12^. However, excessive iron could also drive ROS production^13^. Since the inhibition of complex I by rotenone only induced ROS burst in bat, but not human, cells, we reasoned that bat cells may have higher level of iron to stimulate ROS production. We improved the bathophenanthroline based assay^14,15^ to unambiguously measure the levels of both ferrous (Fe^2+^) and ferric (Fe^3+^) ions in the same sample (Figure 2C). Contrary to our expectation, bat cells exhibited lower level of Fe^2+^ and similar level of Fe^3+^ (thus lower total iron content) comparing to human cells (Figure 2D). Therefore, the ROS burst triggered specifically in bat, but not human, cells upon rotenone treatment was not due to more iron content in bat cells.

**Figure 2.**
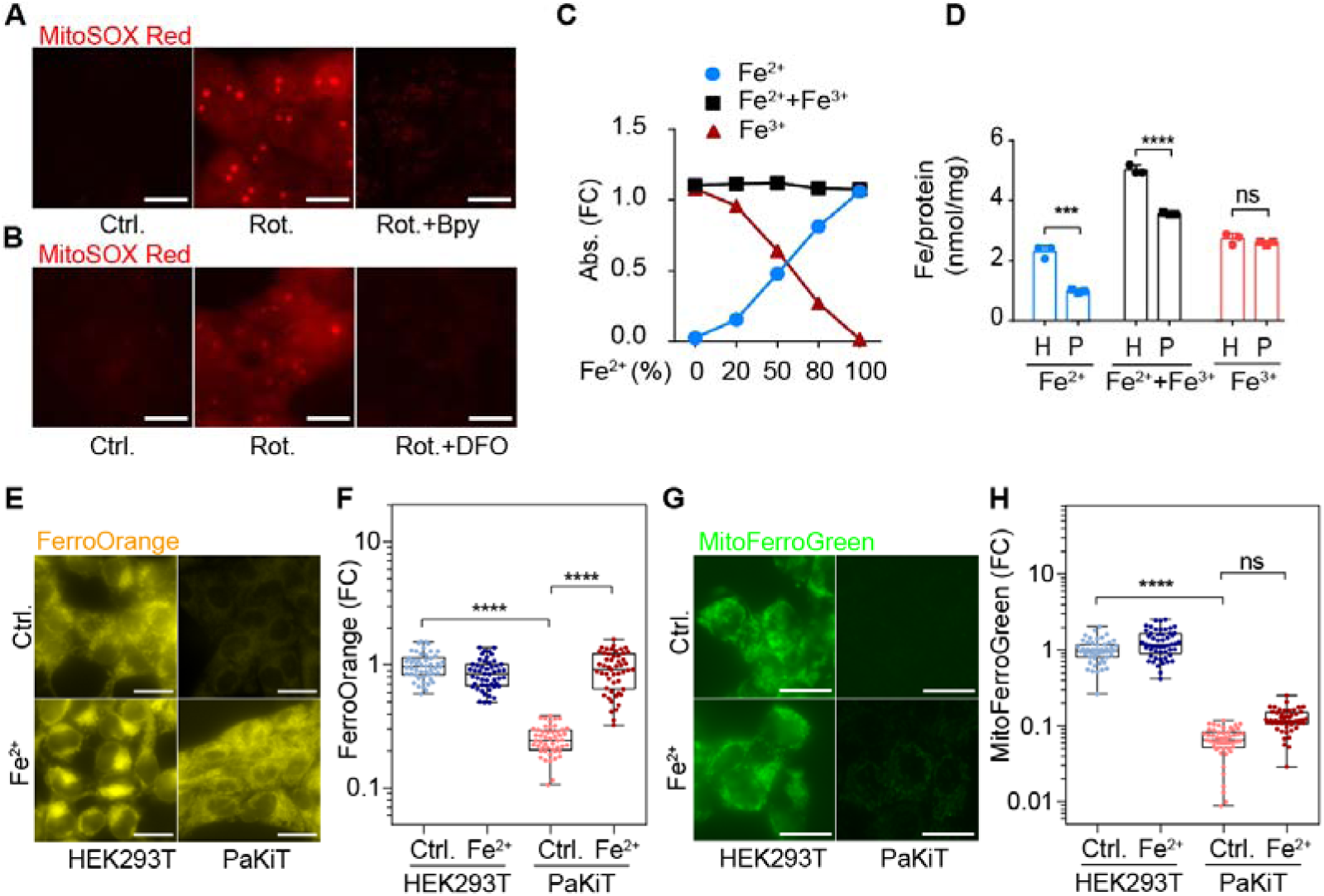
Iron homeostasis is required for mitochondrial ROS burst in bat cells. (A) PaKiT cells were pre-treated with 100 μM 2,2’-bipyridine (Bpy) for 1 day followed by rotenone (Rot.) treatment (50 μM) for 4 hours before live cell imaging of ROS signal. (B) PaKiT cells were pre-treated with 500 μM deferoxamine (DFO) for 1 day before receiving 50 μM rotenone for 4 hours and imaging for ROS signal. (C) An improved bathophenanthroline-based colorimetric assay for mixed ammonium ferrous (Fe^2+^) sulfate and ammonium ferric (Fe^3+^) citrate at indicated ratio (500 μM total iron). Abs.: absorption. FC: fold change. (D) Iron content analysis of HEK293T (H) and PaKiT (P) by bathophenanthroline-based colorimetric assay (n = 3, mean ± s.d.). (E) HEK293T and PaKiT cells were supplemented with 100 μM ferrous ion (Fe^2+^) for 1 day before live cell imaging of FerroOrange signal. (F) Quantification of (E). (G) Cells were treated as in (E) before imaging of MitoFerroGreen signal. (H) Quantification of (G). n > 49 cells for each group in (F and H). Statistical analysis was performed using two-tailed Student’s *t*-test for (D), one-way ANOVA with Tukey’s test for multiple comparisons for (F and H), \*\*\**P* < 0.001, \*\*\*\**P* < 0.0001, ns: not significant. Scale bar = 20 μm. See also Figure S2.

To further analyze the iron content in intact cells, labile Fe^2+^ was monitored by FerroOrange in living cells. Consistent with our biochemical measurement, the level of Fe^2+^ was significantly lower in bat cells comparing to human cells (Figure 2E and 2F). Of note, supplementing Fe^2+^ in culture medium significantly increased the FerroOrange signal in bat cells (Figure 2E and 2F). To specifically monitor the mitochondrial Fe^2+^ pool, living cells were stained by MitoFerroGreen. Again, the level of mitochondrial Fe^2+^ in bat cells was significantly lower comparing to human cells (Figure 2G and 2H). Importantly, supplementing Fe^2+^ in culture medium had marginal effect on bat mitochondrial iron content (Figure 2G and 2H), indicating that the mitochondrial iron import is tightly controlled and separately regulated from the bulk iron content (measured by FerroOrange) within bat cells. Of note, chelating Fe^2+^ by Bpy further diminished the signal of both FerroOrange and MitoFerroGreen (Figure S2C-S2F).

### Complex I inhibition triggers mitochondrial iron overload

To investigate how iron homeostasis may be regulated upon complex I inhibition, living cells were stained by FerroOrange or MitoFerroGreen after rotenone treatment. Surprisingly, complex I inhibition led to an obvious decrease of the bulk Fe^2+^ level as well as a concomitant increase of the mitochondrial Fe^2+^ content in bat cells (Figure 3A-3D. Thus, complex I serves as a trigger to regulate mitochondrial iron import within bat cells. To further analyze whether mitochondrial iron overload is upstream of ROS burst, complex I was inhibited with rotenone for different time periods. Mitochondrial iron signal was significantly elevated within half hour, while mitochondrial ROS burst did not happen yet (Figure 3E-3G). Upon prolonged inhibition, the ROS burst signal quickly reached plateau within one hour (Figure S3A-S3D). Together, these data demonstrated that complex I inhibition triggers mitochondrial iron overload for ROS burst.

**Figure 3.**
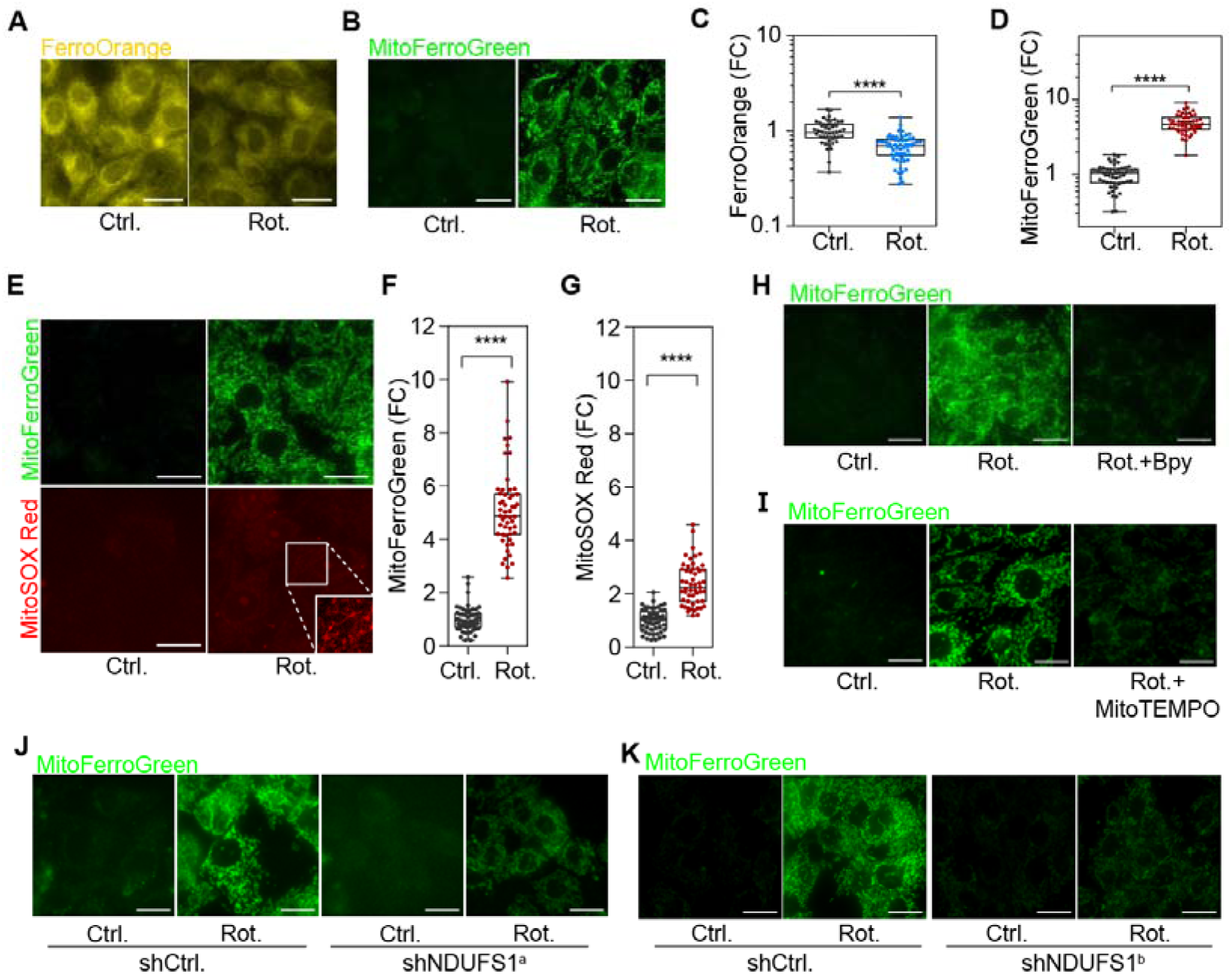
Complex I inhibition triggers mitochondrial iron overload for ROS burst. (A and B) PaKiT cells were treated with 50 μM rotenone (Rot.) for 6 hours before imaging of FerroOrange (A) or MitoFerroGreen (B). (C and D) Quantification of (A and B). n > 49 cells in each group. FC: fold change. (E) PaKiT cells were treated with 10 μM rotenone for 30 min before imaging of MitoFerroGreen or MitoSOX Red. The white squared area was zoomed in with adjusted contrast for clarity. (F and G) Quantification of (E). n > 52 cells in each group. (H) PaKiT cells were treated with 100 μM 2,2’-bipyridine (Bpy) for 1 day before receiving 50 μM rotenone for 1 hour and imaging of MitoFerroGreen. (I) PaKiT cells were pre-treated with 100 μM MitoTEMPO for 1 day before receiving 50 μM rotenone for 1 hour and imaging of MitoFerroGreen. (J and K) NDUFS1 was knocked down by shRNA for 2-3 days in PaKiT before 50 μM rotenone treatment for 1 hour and imaging of MitoFerroGreen. Statistical analysis was performed using two-tailed Student’s *t*-test for (C, D, F and G), \*\*\*\**P* < 0.0001. Scale bar = 20 μm. See also Figure S3.

As expected, the iron chelator Bpy inhibited the rotenone-induced MitoFerroGreen signal, demonstrating the specificity of iron detection (Figure 3H). Interestingly, the ROS scavenger MitoTEMPO effectively suppressed the rotenone-induced mitochondrial iron level (Figure 3I). Together with the observation that mitochondrial iron overload is upstream of ROS burst (Figure 3E-3G), these data indicated that the initial ROS produced by complex I inhibition is required to stimulate the mitochondrial iron overload, which in turn amplifies the ROS signal and eventually leads to ROS burst. To assess whether the rotenone-induced mitochondrial iron overload depended on complex I, the expression level of NDUFS1 was knocked down by shRNA. The mitochondrial iron overload was effectively inhibited by knocking down NDUFS1 expression, supporting the specific involvement of complex I in the regulation of iron translocation upon rotenone treatment (Figure 3J and 3K). Taken together, we conclude that complex I inhibition triggers mitochondrial iron overload to stimulate ROS burst in bat cells.

### Mitochondrial iron influx requires **ΔΨ**m and MCU

The ΔΨm has been reported to be required for iron import into the matrix of yeast mitochondria^16^. During the course of iron content analysis, we noticed that the mitochondrial iron overload was accompanied by elevated ΔΨm in bat cells (Figure S4A and S4B). To address whether the ΔΨm was required for the rotenone-induced iron overload into bat mitochondria, the proton ionophore carbonyl cyanide m-chlorophenyl hydrazone (CCCP) was utilized to dissipate the ΔΨm (Figure 4A). Remarkably, the rotenone-induced signal of MitoFerroGreen was not enriched in mitochondria in the presence of CCCP, but evenly distributed within bat cells (Figure 4B), indicating that the ΔΨm was crucial for mitochondrial iron import in bat cells. Indeed, when both complexes I and III were inhibited by the combination of rotenone and antimycin A, the ΔΨm was also effectively suppressed (Figure 4C). A dispersed signal of MitoFerroGreen was again observed upon rotenone treatment in the presence of antimycin A (Figure 4D). Of note, both antimycin A or CCCP treatment impaired rotenone-induced ROS signal (Figure S4C and S4D). Together, we conclude that the mitochondrial iron influx induced by rotenone depends on the ΔΨm in bat cells.

**Figure 4.**
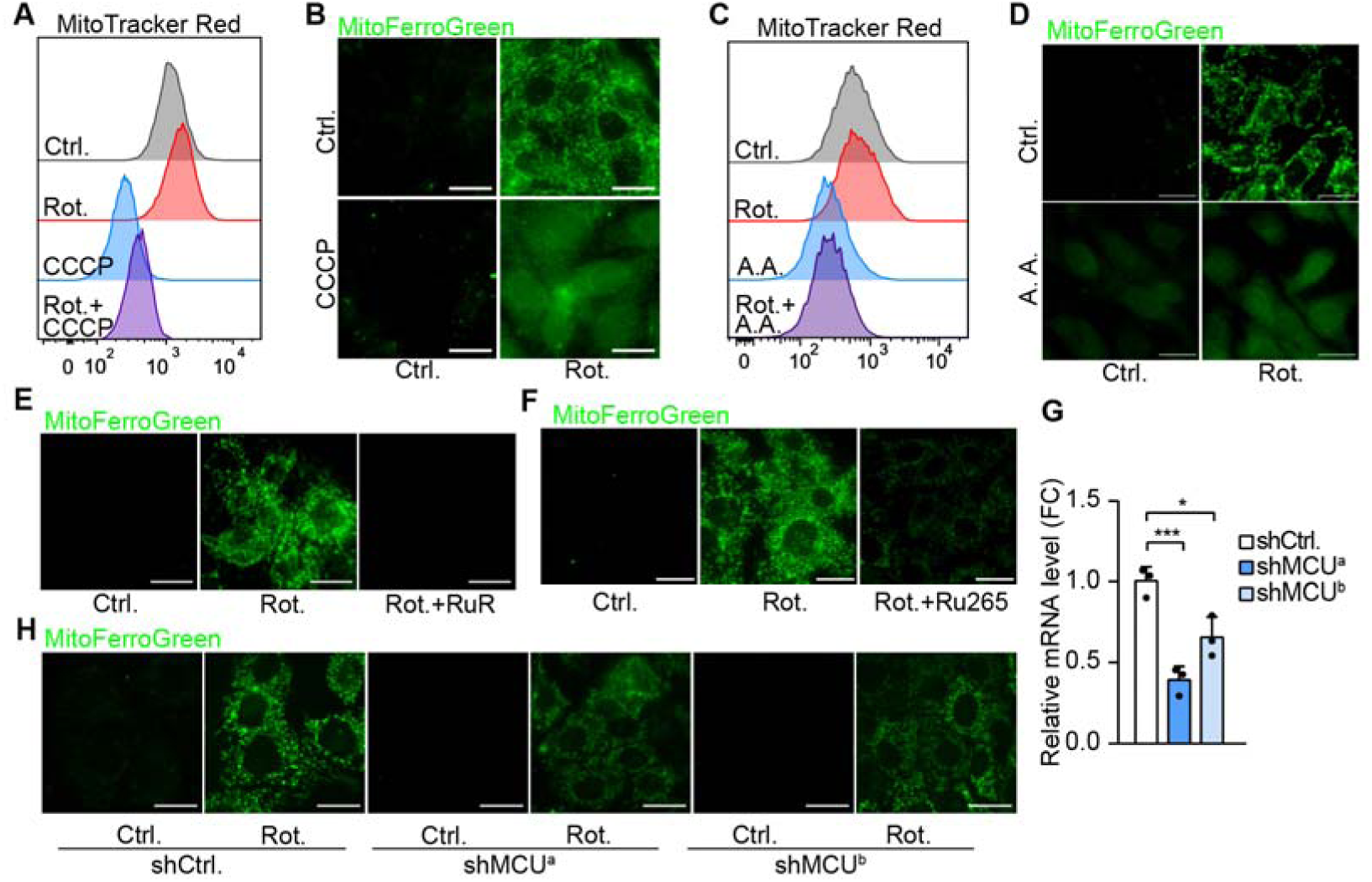
Mitochondrial iron influx depends on **ΔΨ**m and MCU. (A) PaKiT cells were treated with 50 μM rotenone or/and 10 μM CCCP as indicated for 1 hour before flow cytometry analysis of ΔΨm (MitoTracker Red). (B) PaKiT cells were treated as in (A) before imaging of MitoFerroGreen. (C) PaKiT cells were treated with 50 μM rotenone or/and 50 μM antimycin A (A.A.) as indicated for 1 hour before flow cytometry analysis of ΔΨm. (D) PaKiT cells were treated as in (C) before imaging of MitoFerroGreen. (E) PaKiT cells were pre-treated with 50 μM ruthenium red (RuR) for 1 day before rotenone treatment (50 μM) for 1 hour and imaging of MitoFerroGreen. (F) PaKiT cells were pre-treated with 100 μM Ru265 for 1 day before rotenone treatment (50 μM) for 1 hour and imaging of MitoFerroGreen. (G) RT-qPCR analysis of MCU expression (normalized to β-Actin) in PaKiT cells expressing two different shRNAs against MCU for 3 days (n = 3, mean ± s.d.). (H) MCU was knocked down by two different shRNAs for 3 days before rotenone treatment (10 μM) for 1 hour and imaging of MitoFerroGreen. Statistical analysis was performed using one-way ANOVA with Dunnett’s test for multiple comparisons for (G), \**P* < 0.05, \*\*\**P* < 0.001. Scale bar = 20 μm. See also Figure S4.

To further explore the molecular components regulating mitochondrial iron overload in bat cells, RNA-sequencing (RNA-seq) experiment was performed. The Gene Ontology (GO) enrichment analysis revealed that genes involved in mitochondrial transport were differentially regulated in bat cells upon rotenone treatment (Figure S4E, Table S3). Interestingly, in the mitochondrial transport category, we noticed that genes involved in calcium homeostasis (including MCU) were enriched (Figure S4F). However, the expression level of reported mitochondrial iron transporters (MFRN1/2) was not changed significantly (Figure S4F)^17,18^. To understand the function of MCU in mitochondrial iron overload, ruthenium red (RuR) or Ru265 was utilized to inhibit MCU activity in bat cells treated with rotenone^19^. Remarkably, mitochondrial iron overload and the ROS burst were effectively suppressed by inhibiting MCU (Figure 4E, 4F and S4G). Importantly, knocking down MCU expression by two different shRNAs also impaired rotenone-induced mitochondrial iron overload and ROS burst (Figure 4G, 4H and S4H). Taken together, we conclude that complex I inhibition triggers mitochondrial iron overload for the ROS burst in MCU and ΔΨm dependent manners.

### VDAC is crucial for mitochondrial iron import

How iron is transported across mitochondrial outer membrane remains elusive. VDAC is the most abundant outer membrane protein mediating the flux of metabolites and ions across mitochondrial outer membrane, but whether it participates in mitochondrial iron import is still unknown^20,21^. To explore the possible involvement of VDAC in mitochondrial iron overload, the expression of VDAC was knocked down by two different shRNAs (Figure S5A and S5B). Importantly, the downregulation of VDAC expression effectively suppressed rotenone-induced mitochondrial iron overload and ROS burst (Figure S5C-S5E). These data indicated that VDAC was required for mitochondrial iron overload upon complex I inhibition.

The oligomeric states of VDAC play important roles in the regulation of outer membrane permeability^22^. To obtain further insight regarding how VDAC oligomerization may impact mitochondrial iron import, VBIT-4, a potent VDAC oligomerization inhibitor^22,23^ , was added to bat cells receiving rotenone. Surprisingly, VBIT-4 treatment further enhanced rotenone-induced mitochondrial iron overload and dramatically increased ROS production (Figure 5A-5D). This observation indicated that inhibiting VDAC oligomerization may promote mitochondrial iron overload. Towards this end, we treated bat cells with VBIT-4 alone and indeed observed that inhibiting VDAC oligomerization decreased the bulk Fe^2+^ level with a concomitant increase of the mitochondrial Fe^2+^ content (Figure 5E-5H). Time course analysis confirmed that mitochondrial iron overload preceded ROS burst (Figure 5I and 5J). Taken together, these results suggest that VDAC is crucial for rotenone-induced mitochondrial iron import in bat cells.

**Figure 5.**
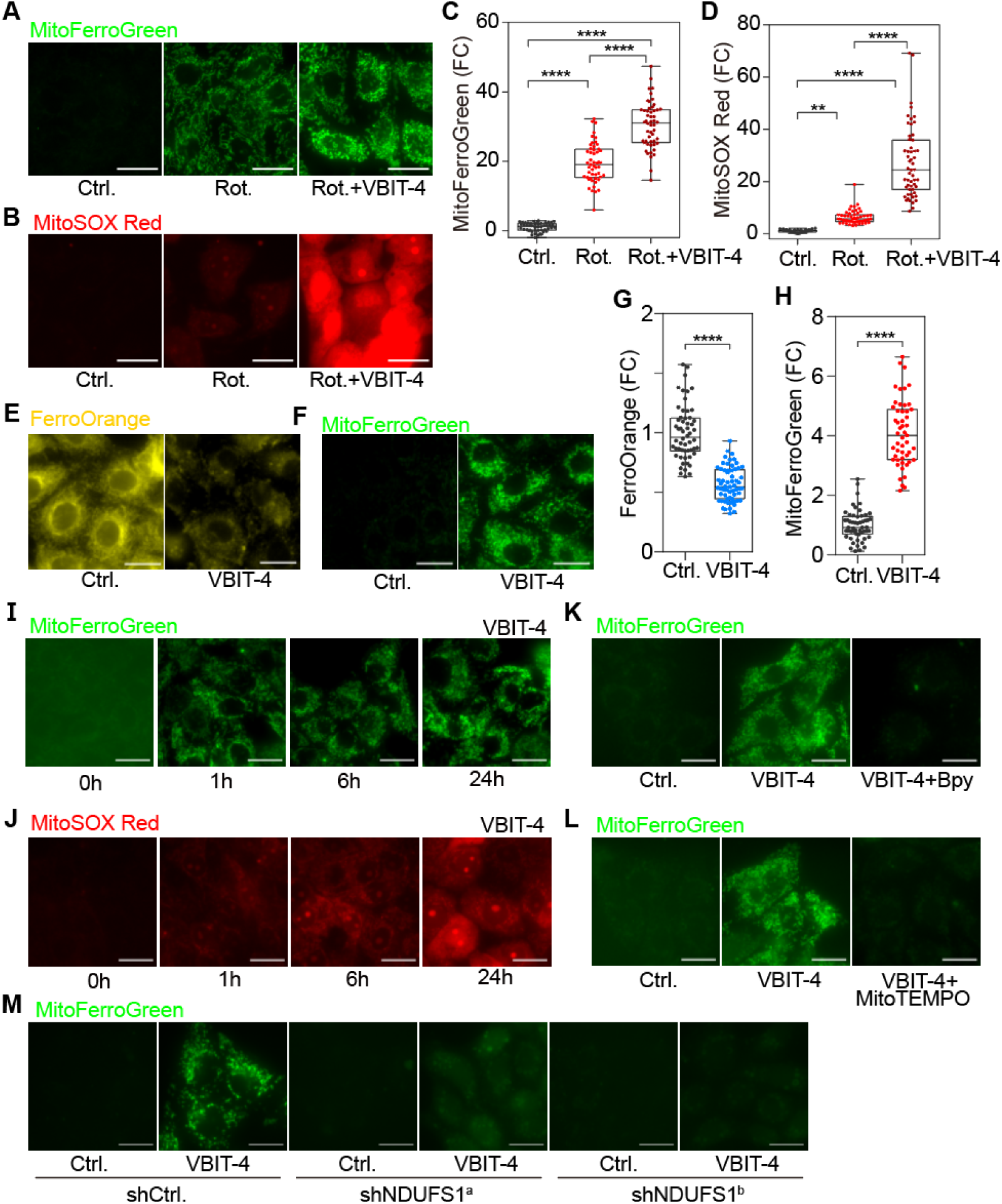
VDAC oligomerization regulates mitochondrial iron import. (A) PaKiT cells were pre-treated with 10 μM VBIT-4 for 1 day before rotenone treatment (50 μM) for 1 hour and imaging of MitoFerroGreen. (B) PaKiT cells were pre-treated with 10 μM VBIT-4 for 1 day before rotenone treatment (50 μM) for 5 hours and imaging of MitoSOX Red. (C) Quantification of (A). n > 49. (D) Quantification of (B). n > 51. (E and F) PaKiT cells were treated with 20 μM VBIT-4 for 1 day before imaging for FerroOrange (E) or MitoFerroGreen (F). (G and H) Quantification of (E and F) respectively. n > 50. (I and J) PaKiT cells were treated with 20 μM VBIT-4 for indicated time before imaging of MitoFerroGreen (I) or MitoSOX Red (J). (K) PaKiT cells were pre-treated with 100 μM 2,2’-bipyridine (Bpy) for 1 day before receiving 20 μM VBIT-4 for 30 min and imaging of MitoFerroGreen. (L) PaKiT cells were pre-treated with 100 μM MitoTEMPO for 1 day before receiving 20 μM VBIT-4 for 30 min and imaging of MitoFerroGreen. (M) NDUFS1 was knocked down by shRNA for 2 days in PaKiT before receiving 20 μM VBIT-4 for 30 min and imaging of MitoFerroGreen. Statistical analysis was performed using one-way ANOVA with Tukey’s test for (C and D) for multiple comparisons, two-tailed Student’s *t*-test for (G and H), \*\**P* < 0.01, \*\*\*\**P* < 0.0001. Scale bar = 20 μm. See also Figure S5 and Figure S6.

### The regulation of VDAC-gated mitochondrial iron import

To analyze how VDAC-gated mitochondrial iron import is regulated, we first wanted to confirm the iron specificity and VDAC dependence of VBIT-4 induced MitoFerroGreen signal. Indeed, both the iron chelator Bpy and VDAC shRNA could effectively impair mitochondrial iron signal upon VBIT-4 treatment (Figure 5K and S5F). Further, when mitochondrial membrane potential was dissipated by CCCP, VBIT-4 induced iron signal was not enriched in mitochondria (Figure S5G). Moreover, the chemical inhibition of MCU by RuR or Ru265 as well as the genetic inhibition of MCU by two different shRNAs both effectively inhibited VBIT-4 induced mitochondrial iron import (Figure S5H-S5J). Together, these data demonstrated that inhibiting VDAC oligomerization by VBIT-4 induces mitochondrial iron import via VDAC-MCU axis in a ΔΨm dependent manner.

To understand whether VDAC-gated mitochondrial iron import depends on respiratory chain and ROS, we treated bat cells with MitoTEMPO. Notably, inhibiting ROS could effectively suppress VBIT-4 induced mitochondrial iron import (Figure 5L). Moreover, specific depletion of complex I effectively impaired mitochondrial iron signal upon VBIT-4 treatment (Figure 5M). Together, these data indicated that complex I and derived ROS was required to trigger VDAC-gated mitochondrial iron overload. Respiratory complexes can be organized into supercomplexes to facilitate electron transfer and reduce ROS generation^24^. To analyze how mitochondrial iron overload may be connected to respiratory complexes in bat mitochondria, protein complexes were extracted by digitonin and subjected to blue native PAGE (BN-PAGE) followed by immunoblotting. The supercomplexes of I/III/IV were destabilized by rotenone, but not VBIT-4 treatment, demonstrating that the destabilization of respiratory supercomplexes is not necessarily required for mitochondrial iron overload (Figure S6A-S6D).

### Mitochondrial iron overload regulates lipid metabolism

To dissect how mitochondrial iron overload affects mitochondrial metabolism, we first analyzed the transcriptome data obtained from bat cells treated with rotenone or VBIT-4. Among the commonly detected genes, 81 differentially expressed genes were overlapped (fold change above 2 or below its reciprocal, FDR-adjusted *P* value < 0.05). 85% of them (69 genes) were positively correlated (Figure S7A, Table S4). The GO enrichment analysis of the commonly upregulated genes revealed that cellular response to lipid may be affected by mitochondrial iron overload (Figure S7B, Table S4). To gain insight how mitochondrial iron overload may impact lipid metabolism, untargeted metabolomics was performed over bat cells treated with DMSO or rotenone. For the tricarboxylic acid (TCA) cycle, the level of citric acid was dramatically reduced to ∼5% of the control upon rotenone treatment, while the downstream metabolites oxoglutaric acid, fumaric acid and malic acid were significantly upregulated (Figure 6A and 6B). The carbon flow may be sustained by the upregulated level of glutamine and glutamic acid (Figure 6B). Moreover, the NADH/NAD^+^ ratio was increased about four-fold (Figure 6C). Together, these data were consistent with the TCA cycle arrest and impaired energy production upon complex I inhibition. Indeed, the ATP/ADP ratio was reduced by more than half in rotenone treated samples (Figure 6C). Pyruvate can be converted to acetyl-CoA (for citric acid synthesis) or lactate. Interestingly, we noticed that the pyruvate level was reduced by 26%, while the lactate level was increased by 29% (Figure 6D). Therefore, the diversion of pyruvate to lactate can not fully explain the sharp decrease of citric acid upon complex I inhibition.

**Figure 6.**
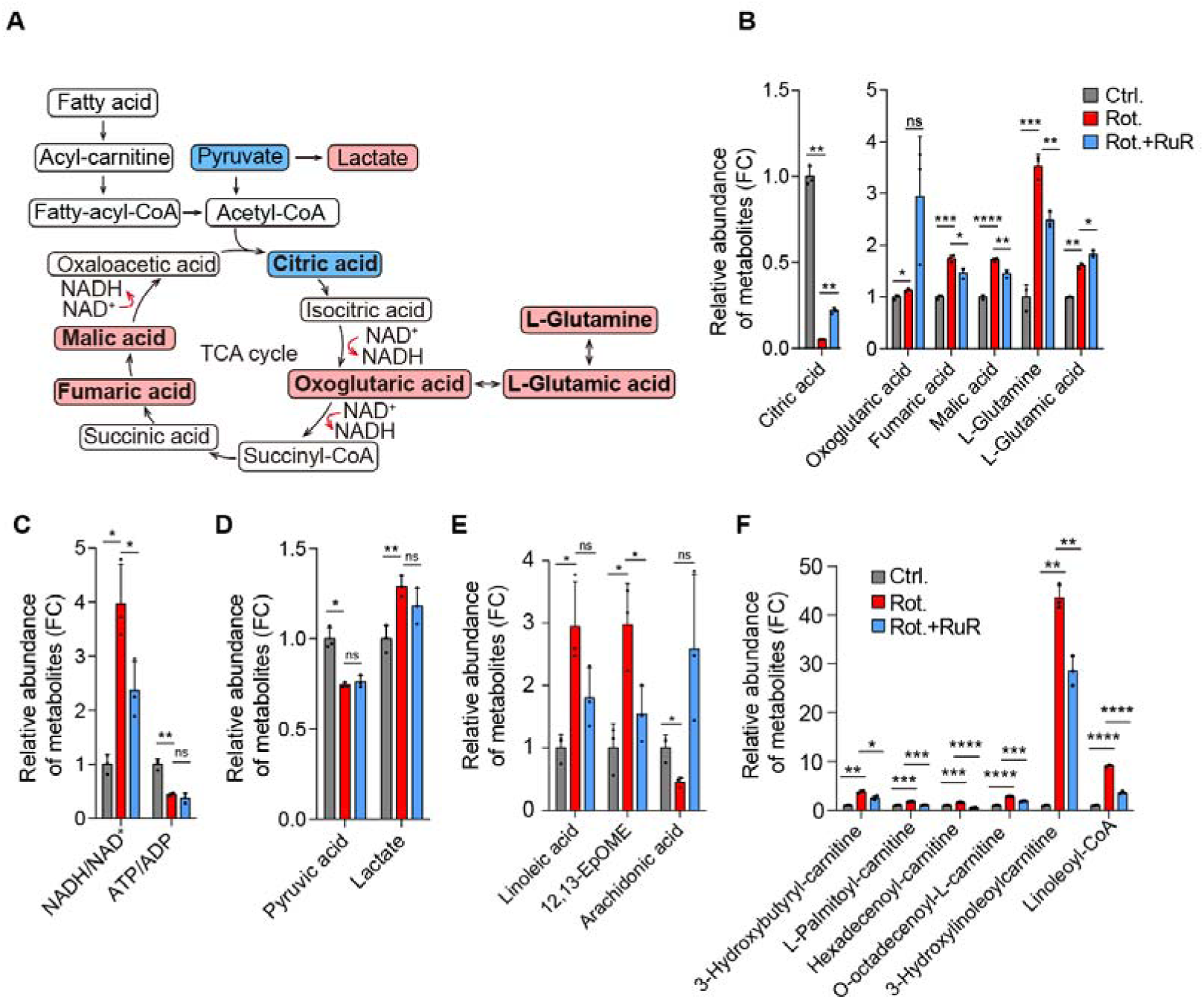
Mitochondrial iron overload remodels lipid metabolism in bat cells. (A) Schematic diagram of the TCA cycle pathway. (B-F) PaKiT cells were pre-treated with 50 μM ruthenium red (RuR) for 1 day before 10 μM rotenone (Rot.) treatment for 1 hour followed by metabolomics analysis. Relative abundance of metabolites involves in TCA cycle (B-D) and lipid metabolism (E-F) (n = 3, mean ± s.d.). Statistical analysis was performed using two-tailed Student’s *t*-test for (B-F), \**P* < 0.05, \*\**P* < 0.01, \*\*\**P* < 0.001, \*\*\*\**P* < 0.0001, ns: not significant. See also Figure S7.

Fatty acid β-oxidation is another important source providing acetyl-CoA for citric acid synthesis (Figure 6A). We reasoned that the observed sharp reduction of citric acid could be due to insufficient acetyl-CoA generated from fatty acid oxidation. Indeed, we observed that linoleic acid and its derived vernolic acid (12,13-EpOME) were accumulated upon complex I inhibition, while arachidonic acid was significantly reduced (Figure 6E and S7C), indicating the conversion of linoleic acid into arachidonic acid was impaired. Moreover, long-chain acylcarnitines were accumulated upon complex I inhibition. Interestingly, we noticed that 3-hydroxylinoleoylcarnitine and linoleoyl-CoA was increased about forty-three-fold and nine-fold, respectively (Figure 6F). Importantly, suppressing mitochondrial iron overload induced by complex I inhibition significantly reduced the level of 3-hydroxylinoleoylcarnitine and linoleoyl-CoA along with other acylcarnitines and concomitantly reversed the ratio of linoleic acid species to arachidonic acid (Figure 6E, 6F and S7C). Of note, the inhibition of ATP production and the diversion of pyruvate to lactate were not affected by reduced mitochondrial iron influx (Figure 6C and 6D). These data demonstrated that fatty acid oxidation was regulated by mitochondrial iron overload.

### Linoleic acid reversibly promotes lipid droplet accumulation

The accumulation of excessive amount of fatty acids can lead to lipid droplet formation. To analyze how mitochondrial iron translocation may impact the level of lipid droplets, bat cells were treated with rotenone or VBIT-4 to induce iron translocation into mitochondria followed by BODIPY staining. Imaging analysis revealed that both the intensity and the size of lipid droplets were significantly elevated upon mitochondrial iron translocation (Figure 7A-7F). Furthermore, washing out rotenone or VBIT-4 for one hour effectively reversed the mitochondrial iron translocation as well as the lipid droplet accumulation (Figure 7G-7L). Since mitochondrial iron translocation upregulated the level of linoleic acid related molecules, we were curious to know whether linoleic acid alone could promote lipid droplet accumulation. We observed that supplying linoleic acid was sufficient to promote the accumulation of lipid droplets without affecting mitochondrial iron levels (Figure 7M and 7N). Moreover, removing linoleic acid quickly reversed the accumulation of lipid droplets (Figure 7O). Taken together, we conclude that mitochondrial iron-induced linoleic acid species can promote lipid droplet accumulation.

**Figure 7.**
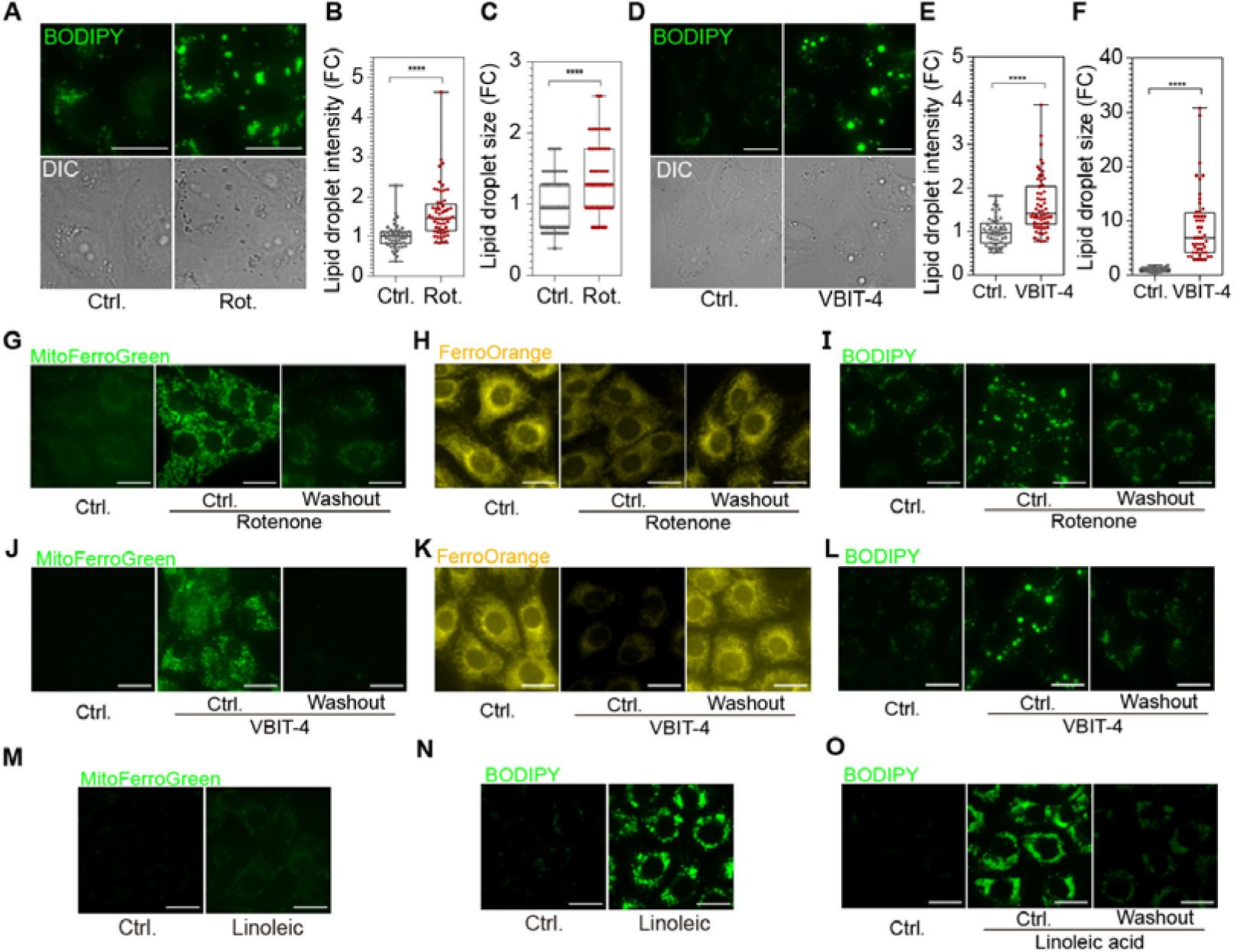
Mitochondrial iron overload promotes lipid droplets accumulation in bat cells. (A and D) PaKiT cells were treated with 50 μM rotenone (Rot.) for 6 hours or 20μM VBIT-4 for 24 hours followed by imaging of BODIPY. (B and C) Quantification of (A). n > 51 for (B), n > 56 for (C). (E and F) Quantification of (D). n > 53 for (E), n > 47 for (F). (G-I) PaKiT cells were treated with 50 μM rotenone for 6 hours and washout for 1 hour followed by imaging of MitoFerroGreen (G), FerroOrange (H) or BODIPY (I). (J-L) PaKiT cells were treated with 20 μM VBIT-4 for 24 hours and washout for 1 hour followed by imaging of MitoFerroGreen (J), FerroOrange (K) or BODIPY (L). (M-N) PaKiT cells were treated with 50 μM of linoleic acid for 24 hours and were imaged using MitoFerroGreen (M) or BODIPY (N). (O) PaKiT cells were treated with 100 μM linoleic acid for 24 hours and washout for 1 hour followed by imaging of BODIPY. Statistical analysis was performed using two-tailed Student’s *t*-test for (B, C, E and F). Scale bar = 20 μm.

## DISCUSSION

As the only mammal capable of powered flight, bats represent a fascinating model for understanding the evolutionary adaptations of mitochondrial energy metabolism. Being proposed key to various aspects of bat biology, including longevity and flight, mitochondrial respiration of bats seems to have adapted to the intensive energy demands and minimal ROS production during million years of evolution^3,4,25–28^. Limited ROS generation may be regulated at the level of complex I in long-lived species, including bats^27,29^. In this study, we uncovered that complex I inhibition triggered mitochondrial iron translocation and subsequent ROS burst, which underscores the crucial role of complex I in regulating cellular iron homeostasis beyond its canonical function as a NADH:ubiquinone oxidoreductase. The mitochondrial iron overload depended on the ΔΨm in bat cells as reported in other model organisms^16,30^, since depleting ΔΨm by CCCP or by the combined inhibition of complexes I and III impaired the mitochondrial enrichment of iron.

Different inner membrane proteins with potential redundancy are reported for iron transport across the mitochondrial inner membrane in various experimental models^17,31,32^. We discovered that MCU facilitated mitochondrial iron import in bat cells. Interestingly, MCU and calcium are reported to contribute to the mitochondrial ROS production in other experimental models^33–35^. Whether and how the homeostasis of calcium and iron is coupled in bats as well as in other organisms is worthy of further investigation. As the most abundant mitochondrial outer membrane protein, the β-barrel protein VDAC has been reported to assemble into various forms of oligomers^36–38^. The channel activity of VDAC is voltage-gated with anion preferred in its open state and cation selective in its closed state^39^. Although VDAC and other proteins have been proposed for iron transport across mitochondrial outer membrane, experimental evidence is scarce^20^. We revealed that suppressing VDAC expression impaired mitochondrial iron overload. VDAC exists in different isoforms and various oligomeric states. The oligomerization of VDAC has been reported to be required for the release of mitochondrial contents into cytosol. Importantly, inhibiting VDAC oligomerization by VBIT-4 promoted mitochondrial iron import in bat cells. Intriguingly, the VDAC-gated mitochondrial iron import was complex I dependent. It is currently unknown how the activity of complex I is interconnected to the oligomeric status of VDAC. But possible interaction of VDAC and complex I has been proposed^40^. Since mitochondrial ROS is involved in the signal transmission for iron import, it is worth to further investigate whether proteins of cellular iron storage, mobilization and transport systems are subjected to redox regulation in bat cells.

Iron serves essential roles in mitochondrial energy metabolism by being a key component of enzymes involved in fatty acid β-oxidation, the TCA cycle and the OXPHOS system^10^. Disturbances of iron homeostasis can cause abnormal lipid metabolism, leading to mitochondrial damage and metabolic diseases^41^. We demonstrated that iron translocation into bat mitochondria governed by complex I disrupted fatty acid oxidation and led to the significant upregulation of long-chain acylcarnitines as well as linoleic acid derivatives. This further resulted the reversible accumulation of lipid droplets, which appeared to serve as temporary depots to store excessive lipids. The accumulation of linoleic acid derivatives was downstream of mitochondrial iron overload, thus could be alleviated by reversing mitochondrial iron level. This was further supported by the observation that supplementing linoleic acid alone was sufficient to drive lipid droplet accumulation without affecting mitochondrial iron level.

In summary, we propose that complex I functions as a central regulatory hub governing mitochondrial iron translocation to coordinate respiration with lipid metabolism (Figure S8). This study provides the first evidence that the precise regulation of mitochondrial iron levels may be pivotal to the evolutionary adaptation of mitochondrial energy metabolism in bat cells. It would be intriguing to investigate whether the mechanism identified in this study play roles in the metabolic adaptation of bats in their natural habitats.

## ACKNOWLEDGEMENTS

We thank Dr. Nils Wiedemann, Dr. Xiaorong Zhang, Dr. Hongbo Zhang and the members of the Qiu lab for discussion and critical comments on the manuscript. We thank Zhigang Peng, Biyue Tian and Tiansheng Chou for technique support. We thank Dr. Peng Zhou from Wuhan Institute of Virology, Chinese Academy of Sciences for sharing PaKiT and HEK293T cells. This work was supported by grants from the National Natural Science Foundation of China (32270722, 82230023, 82400873), the Natural Science Foundation of Hunan Province (2022JJ30914, 2025JJ60551), Postdoctoral Fellowship Program (Grade B) of China Postdoctoral Science Foundation (GZB20240866), Central South University (2023QYJC035) and Feifan Scholar Fund of Xiangya Hospital of Central South University.

## AUTHOR CONTRIBUTIONS

J.Q. designed the research and wrote the paper. W.Q.Z. and P.W.P. performed most of the experiments and wrote part of the paper. D.D.W. and L.L.C. performed RNAi experiments and data processing. W.Q.Z. and Z.W. analyzed RNAseq data. C.H.C. and Y.X. performed metabolomics experiments and data analysis. All authors contributed to the analysis and discussion of the results of the experiments.

## SUPPLEMENTAL INFORMATION

Document S1. Figures S1–S8 and Tables S1–S2

Tables S3–S4. GO enrichment analysis of differentially expressed genes.

Supplementary Information is available for this paper. Correspondence and requests for materials should be addressed to J.Q.

