## Supplementary Figures&Methods for "Complex I Governs Iron Levels in Bat Mitochondria to Couple Respiration with Lipid Metabolism"

**Affiliations:**

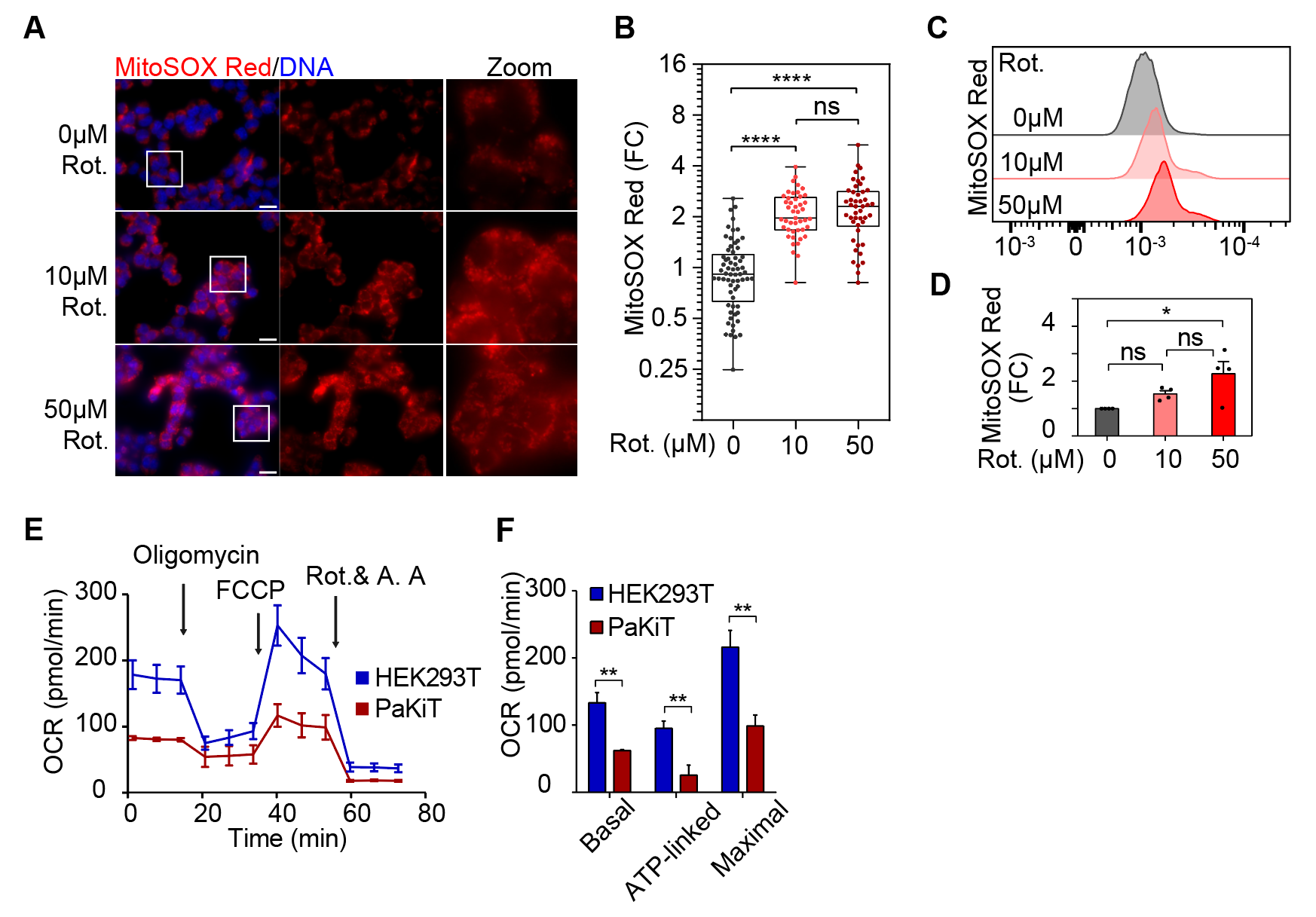

**Figure S1. Complex I inhibition induces mitochondrial ROS burst in bat, but not human, cells, related to Figure 1**

(A) HEK293T cells were treated with rotenone (Rot.) at indicated concentration for 6 hours before imaging of MitoSOX Red. The white squared area was zoomed in.

(B) Quantification of (A) for signal intensity. n > 44 cells in each group.

(C) HEK293T cells were treated as in (A) for flow cytometry analysis of MitoSOX Red.

(D) Quantification of (C). n = 4. mean ± s.e.m.

(E and F) The oxygen consumption rate (OCR) measurement of HEK293T and PaKiT cells. A.A.: antimycin A. n = 4, mean ± s.e.m. Statistical analysis was performed using one-way ANOVA with Tukey's test for multiple comparisons for (B and D), two-tailed Student’s *t*-test for (F), **P* < 0.05, ***P* < 0.01, *****P* < 0.0001, ns: not significant. Scale bar = 20 μm.

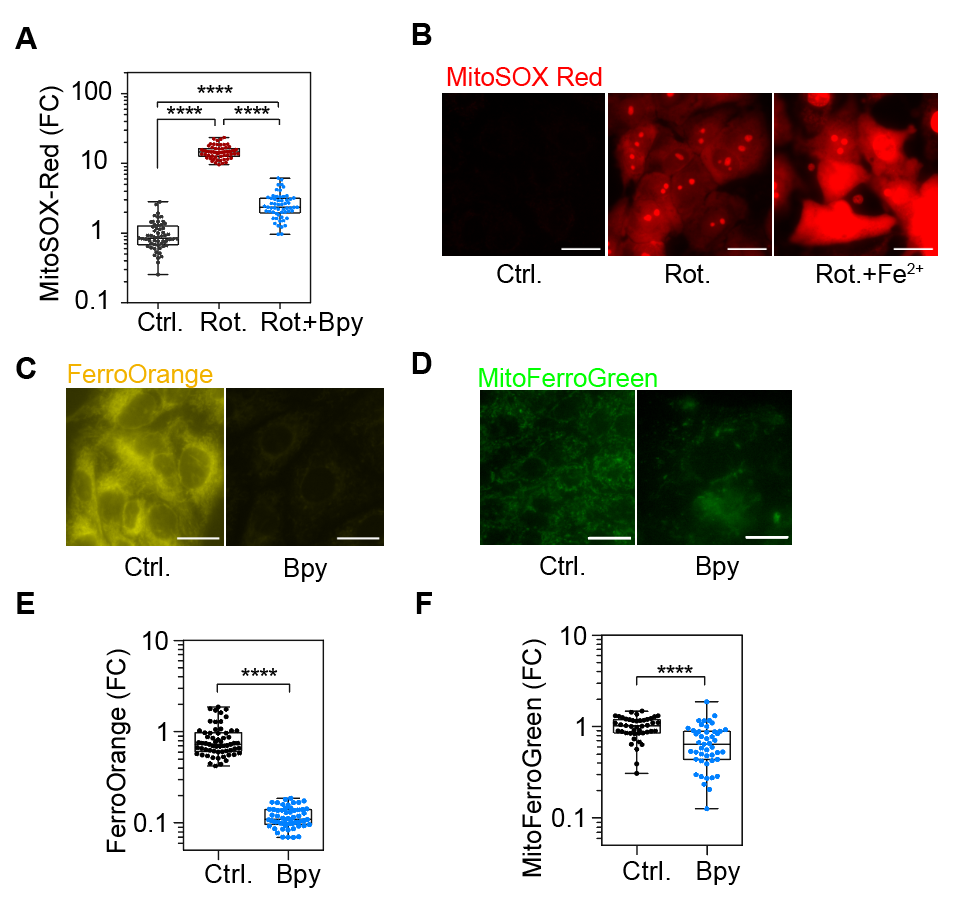

**Figure S2. Iron homeostasis is required for mitochondrial ROS burst in bat cells, related to Figure 2**

(A) Quantification of (Figure 2A). n > 56 cells in each group.

(B) PaKiT cells were pre-treated with 100 μM ammonium ferrous (Fe^2+^) sulfate for 1 day before receiving 50 μM rotenone (Rot.) for 5 hours and imaging for ROS signal.

(C and D) PaKiT cells were treated with 50 μM 2,2'-bipyridine (Bpy) for 1 day before imaging for FerroOrange (C) or MitoFerroGreen signal (D).

(E and F) Quantification of (C) or (D). n > 57 and 43 cells for each group in (C) and (D) respectively. Statistical analysis was performed using one-way ANOVA with Tukey's test for multiple comparisons for (A), two-tailed Student’s *t*-test for (E, F), *****P* < 0.0001. Scale bar = 20 μm.

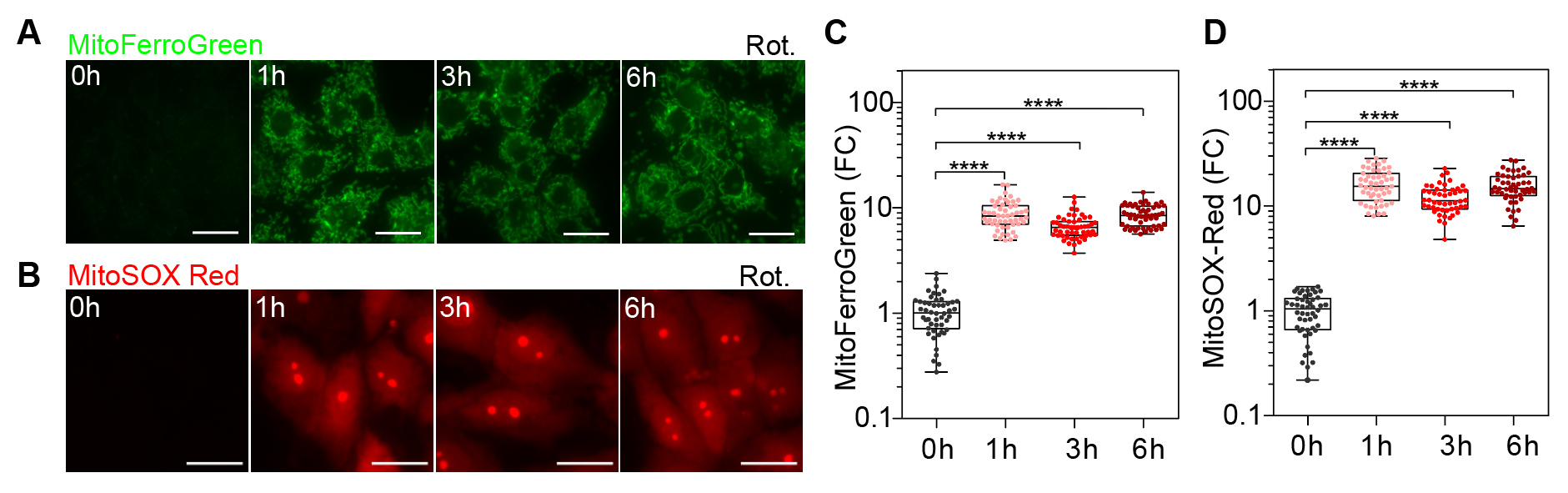

**Figure S3. Complex I inhibition triggers mitochondrial iron overload in bat cells, related to Figure 3**

(A and B) PaKiT cells were treated with 50 μM rotenone for indicated time before imaging of MitoFerroGreen (A) or MitoSOX Red (B).

(C and D) Quantification of (A and B). n > 49 cells in each group. Statistical analysis was performed using one-way ANOVA with Dunnett’s test for multiple comparisons, *****P* < 0.0001. Scale bar = 20 μm.

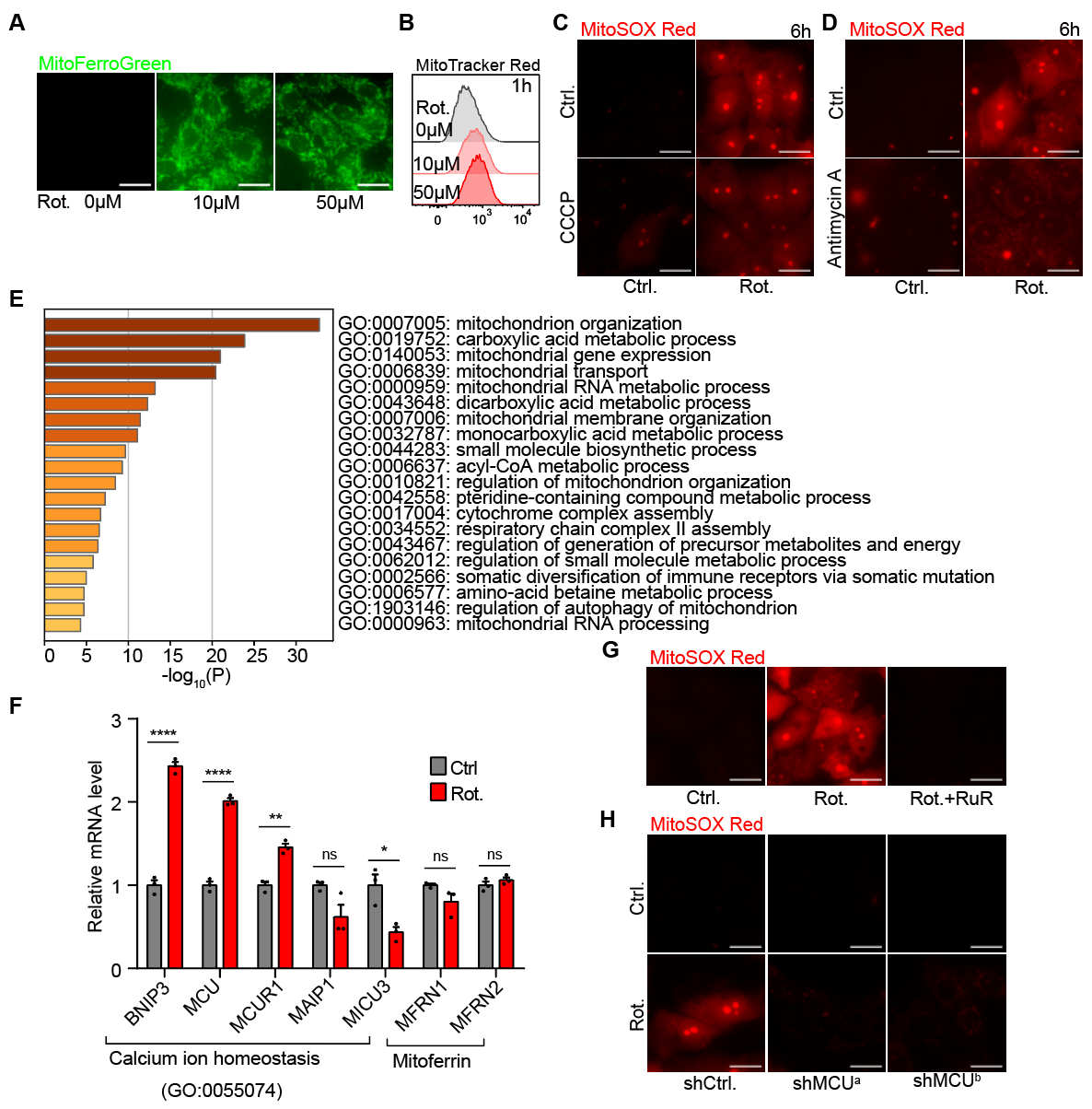

**Figure S4. Mitochondrial iron influx depends on ΔΨm and MCU, related to Figure 4 and Table S3**

(A) PaKiT cells were treated with rotenone (Rot.) at indicated concentration for 1 hour before imaging of MitoFerroGreen.

(B) PaKiT cells were treated as in (A) before flow cytometry analysis of ΔΨm (MitoTracker Red).

(C) PaKiT cells were treated with 50 μM rotenone or/and 10 μM CCCP as indicated for 6 hours before imaging for MitoSOX Red.

(D) PaKiT cells were treated with 50 μM rotenone or/and 50 μM antimycin A (A.A.) as indicated for 6 hours before imaging for MitoSOX Red.

(E and F) PaKiT cells were treated with 50 μM rotenone (Rot.) for 6 hours and analyzed by RNAseq (See Table S3). GO enrichment analysis of 147 differentially expressed mitochondria-related genes (|log2-fold change| > 0.585, *P* < 0.05) using Metascape (E). Relative mRNA expression level of genes involved in calcium ion homeostasis (GO: 0055074) and Mitoferrin were measured by RNAseq (n = 3, mean ± s.e.m.) (F).

(G) PaKiT cells were pre-treated with 50 μM ruthenium red (RuR) for 1 day before rotenone treatment (50 μM) for 6 hours and imaging of MitoSOX Red.

(H) MCU was knocked down by two different shRNAs for 3 days before rotenone treatment (50 μM) for 1 hour and imaging of MitoSOX Red. Statistical analysis was performed using two-tailed Student’s t-test for (F). Scale bar = 20 μm.

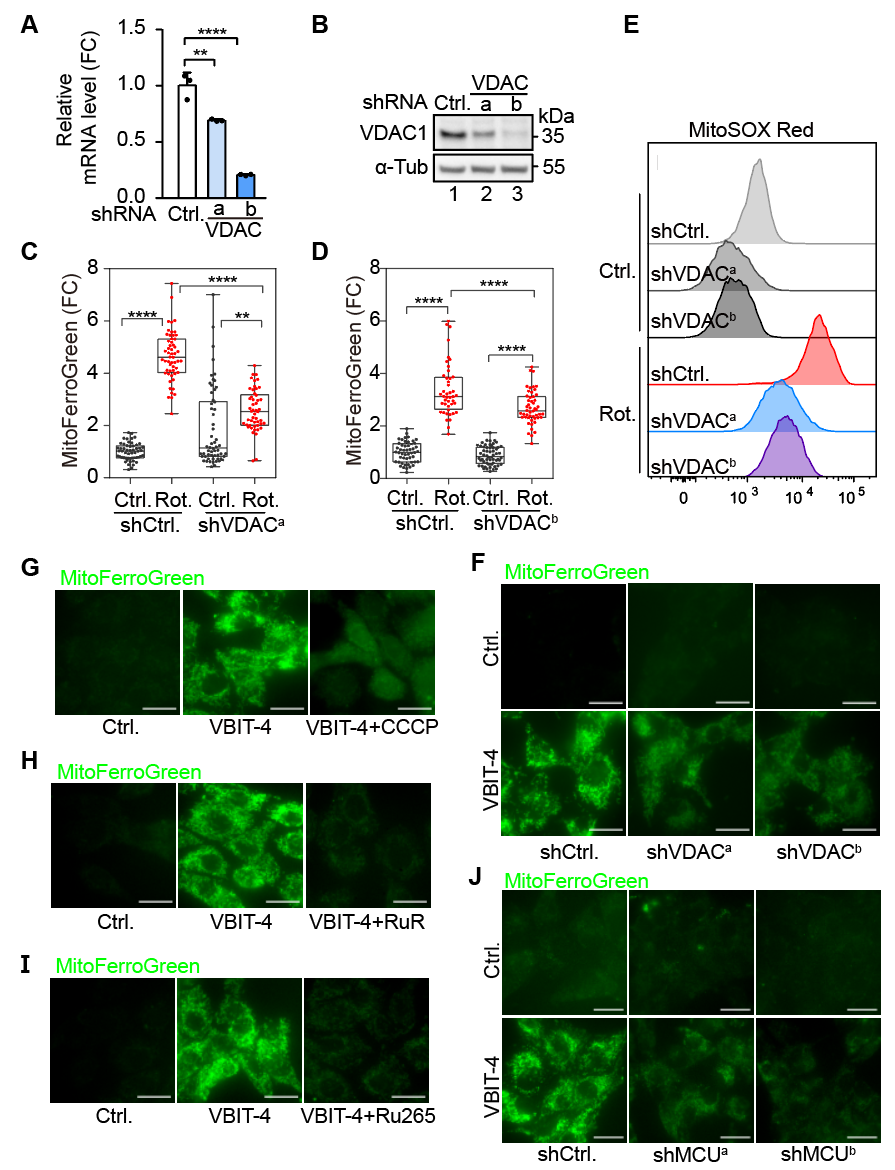

**Figure S5. The regulation of VDAC-gated mitochondrial iron import, related to Figure 5**

(A) VDAC was knocked down by two different shRNAs for 3 days in PaKiT cells before RT-qPCR analysis of VDAC expression (normalized to β-Actin, n = 3, mean ± s.d.).

(B) VDAC was knocked down by shRNA for 2 days in PaKiT cells before SDS–PAGE and immunoblotting with indicated antibodies.

(C-E) VDAC was knocked down by shRNA in PaKiT cells for 3 days before 50 μM rotenone (Rot.) treatment for 1 hour followed by imaging of MitoFerroGreen (C, D) or by flow cytometry analysis of MitoSOX Red (E).

(F) VDAC was knocked down by shRNA for 2 days in PaKiT before receiving 20 μM VBIT-4 for 30 min and imaging of MitoFerroGreen.

(G-I) PaKiT cells were pre-treated with 10 μM CCCP for 0.5h (G), 50 μM RuR for 1 day (H), or 100 μM Ru265 for 1 day (I), before receiving 20 μM VBIT-4 for 30 min and imaging of MitoFerroGreen.

(J) MCU was knocked down by shRNA for 3 days in PaKiT before 20 μM receiving VBIT-4 for 30 min and imaging of MitoFerroGreen. Statistical analysis was performed using one-way ANOVA with Dunnett's test for (A) and with Tukey's test for (C, D) for multiple comparisons, **P < 0.01, ****P < 0.0001. Scale bar = 20 μm.

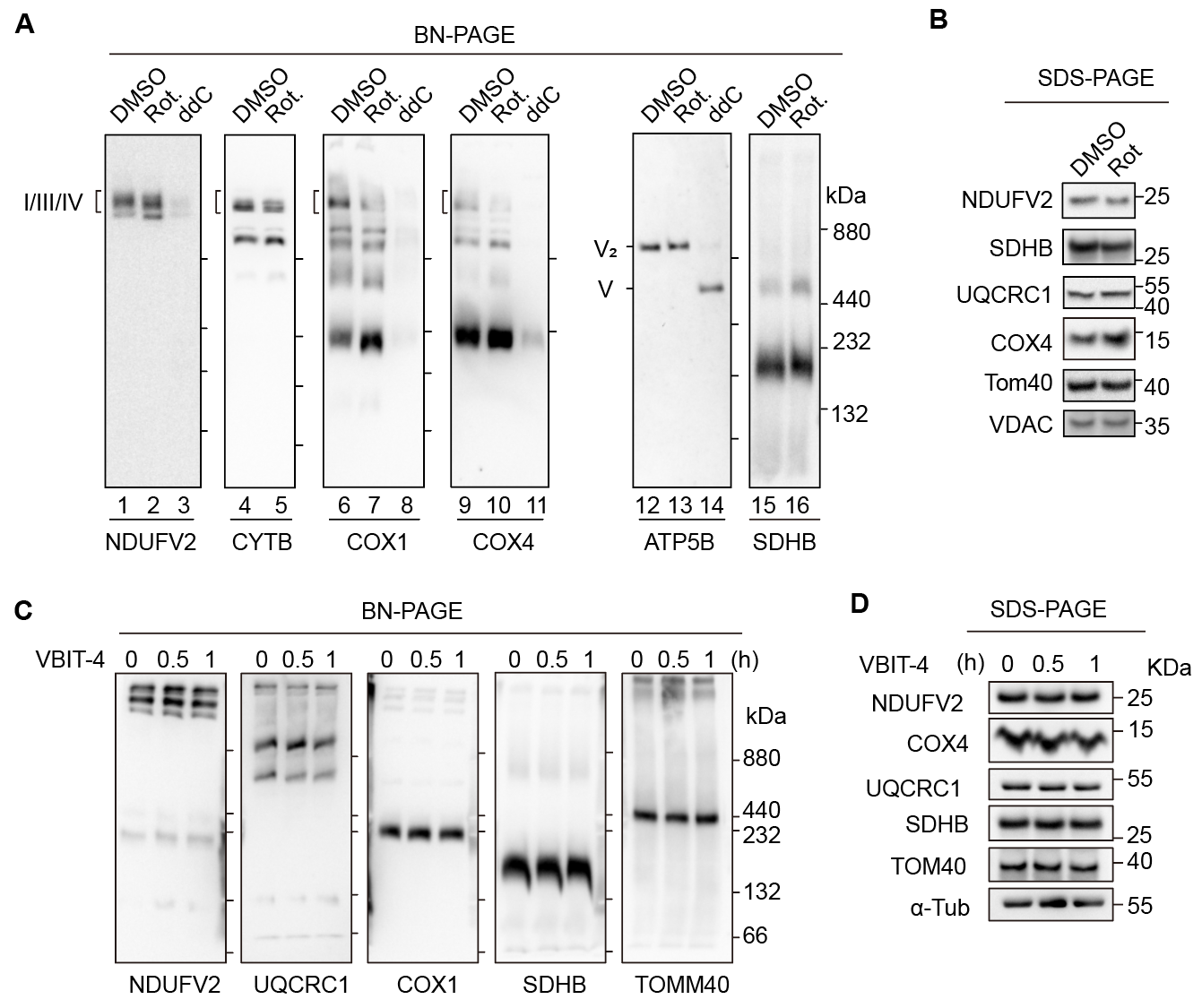

**Figure S6. Inhibiting Complex I, but not VDAC oligomerization, leads to respiratory supercomplexes destabilization in bat cells, related to Figure 5**

(A and B) PaKiT cells were treated with 50 μM rotenone (Rot.) for 1 hour or with 100 μM 2′,3′-dideoxycytidine (ddC) for 5 days before subjected to BN-PAGE (A) or SDS-PAGE (B) and immunoblotting with indicated antibodies.

(C and D) PaKiT cells were treated with 20 μM VBIT-4 for 30 min or 1 hour before subjected to BN-PAGE (C) or SDS-PAGE (D) and immunoblotting with indicated antibodies.

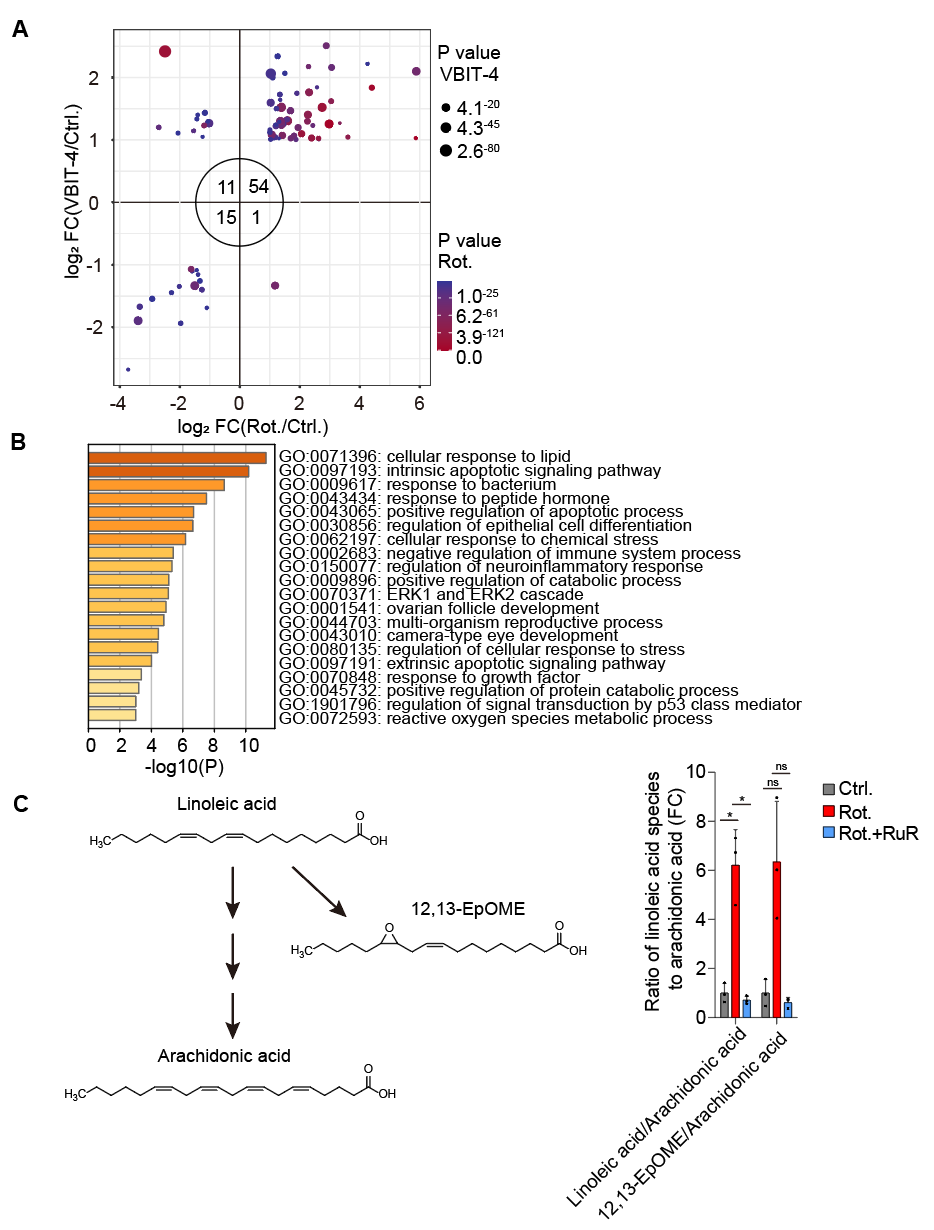

**Figure S7. Mitochondrial iron overload regulates lipid metabolism in bat cells, related to Figure 6 and Table S4**

(A) PaKiT cells were treated with 50 μM rotenone (Rot.) for 6 hours or 20 μM VBIT-4 for 24 h and analyzed by RNAseq. Differentially expressed genes commonly detected in both treatments were plotted. The numbers in the circle represent the number of genes in corresponding quadrant. See Table S4.

(B) The Gene Ontology (GO) enrichment analysis of 54 commonly upregulated genes in (A) using Metascape. See Table S4.

(C) PaKiT cells were pre-treated with 50 μM ruthenium red (RuR) for 1 day before 10 μM rotenone treatment for 1 hour followed by metabolomics analysis. Ratio of linoleic acid species to arachidonic acid was calculated (n = 3, mean ± s.d.). Statistical analysis was performed using two-tailed Student’s *t*-test for (C), **P* < 0.05, ns: not significant.

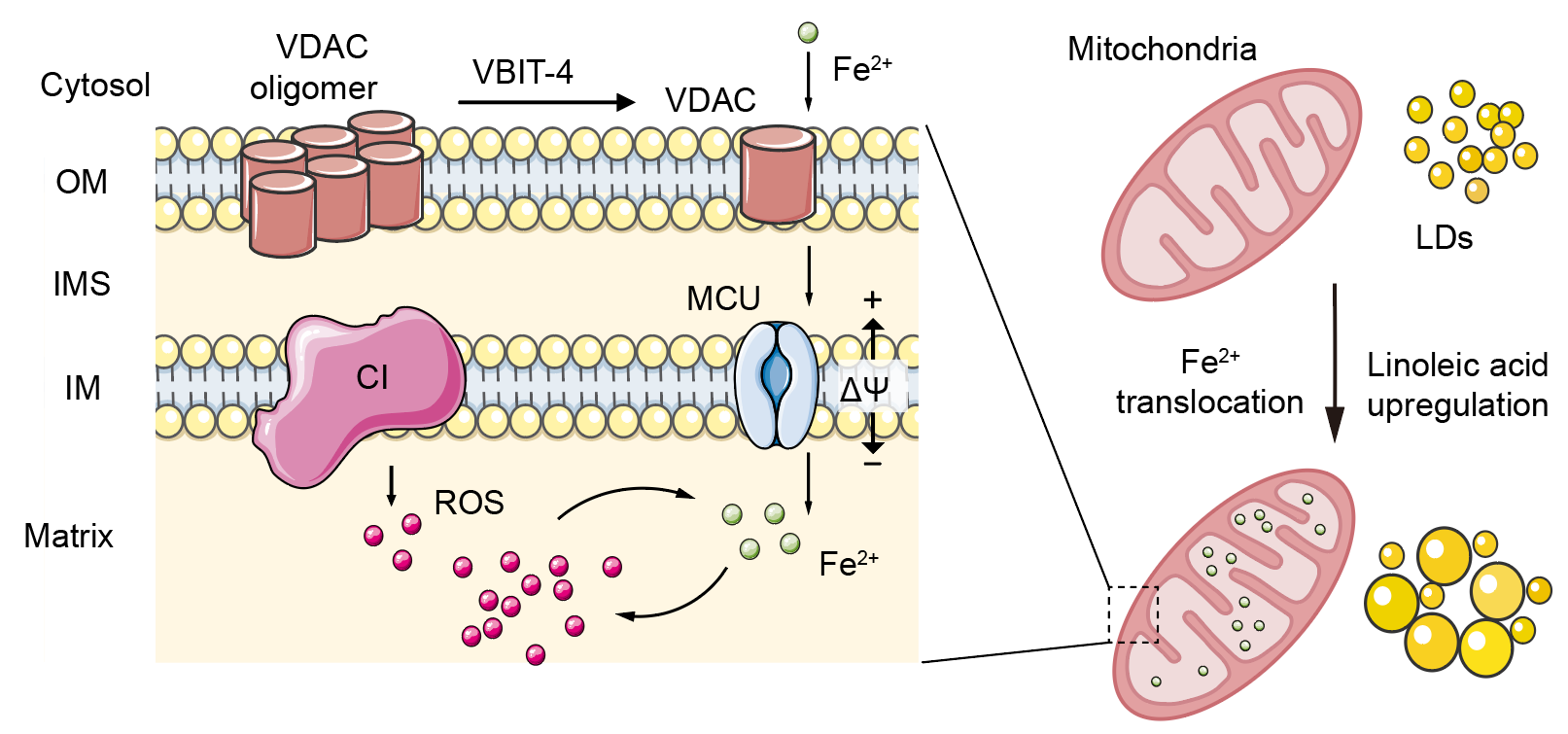

**Figure S8.** **Proposed model.** Complex I functions as a central regulatory hub governing mitochondrial iron translocation via VDAC-MCU axis to coordinate respiration with lipid metabolism.

**Table S1 | Sequences of shRNAs and primers.**

|  | **Name** | **Target sequence (5’ to 3’)** |
| --- | --- | --- |
| shRNAs | shNDUFS1^a^ | GCACAGATTTGCGTTCTAATT |
|  | shNDUFS1^b^ | CTCTCAAAGATTTGCTTAATA |
|  | shMCU^a^ | TAATCAACTCAAGGATGCAAT |
|  | shMCU^b^ | TCAAAGGGCTTAGTGAATATT |
|  | shVDAC^a^ | GGCTATGGATTCGGCTTAATA |
|  | shVDAC^b^ | GCTTGGTCTAGGACTGGAATT |
|  | shCtrl. | CCTAAGGTTAAGTCGCCCTCG |
| Primers  for qPCR | Dloop-F | TCCCCAACTCACGTGAAACC |
|  | Dloop-R | TGTAGGAACCCCCACGGTTTAT |
|  | COX1-F | GGCCCTATCCCAGTATCAAACTC |
|  | COX1-R | TGTGATTCCGGCGGCTAGT |
|  | 18S rRNA-F | CGGCTACCACATCCAAGGA |
|  | 18S rRNA-R | GCTGGAATTACCGCGGCT |
| Primers  for RT-qPCR | MCU-F | TCCACAAGCCAGAGACAGAC |
|  | MCU-R | TGTCGGAGGGGCAGATGTAC |
|  | VDAC1-F | CTGCCTCTCAGAAGATGGCTGT |
|  | VDAC1-R | TTTCCAGACTGCCTGTCACTTT |
|  | β actin-F | CTCCATCATGAAGTGCGATGTTG |
|  | β actin-R | CTTGATCTTCATTGTGCTGGGAG |

**Table S2 | List of antibodies used for immunoblotting.**

| **Antibody (dilution)** | **Company, cat.** |
| --- | --- |
| NDUFS1 (1:1000) | Sangon, D122747 |
| TOM40 (1:1000) | Proteintech, 18409-1-AP |
| α-Tubulin (1:1000) | Sigma, T6199 |
| CYTB (1:1000) | Proteintech, 55090-1-AP |
| VDAC1 (1:1000) | Proteintech, 55259-1-AP |
| VDAC1 (1:1000) | Santa cruz, sc-390996 |
| β-Actin (1:1000) | Sangon, D191047 |
| COX4 (1:1000) | CST, 4850S |
| ATP5B (1:1000) | Santa cruz, sc-55597 |
| SDHB (1:1000) | Proteintech, 10620-1-AP |
| NDUFV2 (1:1000) | Proteintech, 15301-1-AP |
| MTCO1 (1:1000) | Abcam, ab14705 |
| Peroxidase conjugated  Goat Anti-Rabbit IgG (1:10000) | Jackson, 111-035-144 |
| Peroxidase conjugated  Goat Anti-Mouse IgG (1:10000) | Jackson, 115-035-146 |

**STAR Methods**

**Cell culture and generation of stable cell lines.** All cells were cultured in a 37 °C humidified incubator under 5% CO_2_. PaKiT cells were described previously^42^ and obtained from Dr. Peng Zhou from Wuhan Institute of Virology, Chinese Academy of Sciences. PaKiT and HEK293T cells were cultured in DMEM/F-12 (Gibco, C11330500BT) and DMEM (Gibco, C12430500BT), respectively, containing 10% FBS (Procell, 164210-50). Cells were treated with relevant solvent or with rotenone (Sigma, R8875), antimycin A (MKBio, MS0070), CCCP (Sigma, C2759), MitoTEMPO (Santa Cruz, sc-221945A), N-acetyl cysteine (Aladdin, A105422), cyclosporin A (Aladdin, C106893), 2,2'-bipyridine (Aladdin, D108977), deferoxamine (Santa Cruz, sc-203331), ruthenium red (Aladdin, R100832), Ru265 (Sigma, SML2991) or VBIT-4 (Aladdin, V413014) with concentration and period indicated. Scramble shRNA (shCtrl.) or different shRNAs targeting NDUFS1, MCU or VDAC of PaKiT (shRNA sequences were listed in Table S1) were constructed into Tet-pLKO-puro vector using standard molecular cloning methods and verified by sequencing. Plasmids were transfected into HEK293T cells by PEI MAX (Polysciences, 24765) according to the manufacture’s instruction to produce lentiviruses, which were used to infect PaKiT cells (two rounds) supplemented with polybrene (final conc. 8 μg/ml, Sigma, H9268) followed by puromycin selection (3 μg/mL, InvivoGen, ant-pr-1).

**Flow cytometry and live cell imaging.** Cells were stained by 5 μM MitoSOX Red (Thermo, M36008), 100 nM MitoTracker Red (Thermo, M7512), 100 nM MitoTracker Deep Red (Thermo, M22426), 1 μM FerroOrange (Dojindo, F374) or 2 μM MitoFerroGreen (Dojindo, M489) for 30 min at 37 °C. To deplete mtDNA, cells were treated with 100 μM 2′,3′-dideoxycytidine (ddC) (Aladdin, D119514) for 2-5 days. For flow cytometry, cells were trypsin digested (Abiwell, AWC0239a), collected by centrifugation, resuspended in DPBS (Gibco, C14190500BT) and analyzed by BD FACS CantoII or Cytek DxP13 flow cytometers. Data analysis was performed using FlowJo software. For living cell imaging, fluorescent images were acquired by Zeiss Axio Imager.M2 (ApoTome.2) or Axio Observer 7. Special care was taken to avoid oversaturated pixels. Fluorescence intensities were quantified using Fiji software.

**Reverse transcription and quantitative PCR.** To determine mRNA level, total RNA was extracted using TRIzol reagent (Vazyme, R401-01) and measured by NanoDropOneC (Thermo). About 2 μg cDNA was synthesized using RevertAid™ Master Mix (Thermo, M1631). Subsequently, qPCR was then performed directly on the samples using NovoStart® SYBR qPCR SuperMix Plus (Novoprotein, E096-01B) according to the manufacturer's instructions on a QuantStudio 3 System. Primer sequences are listed in Table S1. The results were analyzed using QuantStudio Design and Analysis Software.

**Mitochondrial DNA quantification.** To quantitatively determine the mitochondrial DNA (mtDNA) content, the cells were digested and collected following ddC treatment. They were then lysed with SDS lysis buffer (2% SDS, 20 mM Tris-HCl, pH 7.4, 0.1 mM EDTA, 50 mM NaCl, 10% glycerol) at 95°C for 15 minutes. Subsequently, the proteins in the samples were degraded using Proteinase K (1mg/mL; Coolaber, CP9191) at 37°C for 60 minutes. The enzymatic reaction was terminated by adding PMSF to a final concentration of 1 mM (Aladdin, P105539) and incubating at 95°C for 30 minutes. Subsequently, qPCR was then performed directly on the samples using NovoStart® SYBR qPCR SuperMix Plus (Novoprotein, E096-01B) according to the manufacturer's instructions on a QuantStudio 3 System. Primer sequences are listed in Table S1. The results were analyzed using QuantStudio Design and Analysis Software.

**SDS-PAGE, BN-PAGE and immunoblotting.** For SDS-PAGE, proteins were extracted by SDS lysis buffer (2% SDS, 20 mM Tris-HCl, pH 7.4, 0.1 mM EDTA, 50 mM NaCl, 10% glycerol) supplemented with 1 mM PMSF (Aladdin, P105539) and 1× homemade Protease Inhibitor Cocktail. 200× Protease Inhibitor Cocktail contains 100 mM AEBSF hydrochloride (Macklin, A800781), 0.08 mM Aprotinin (Macklin, A6353), 5 mM Bestatin (Macklin, B802902), 1.5 mM E-64 (Macklin, E808962), 2 mM Leupeptin hemisulfate (Macklin, L812486) and 1 mM Pepstatin (Macklin, P6117). For BN-PAGE, proteins were solubilized with 1% (w/v) digitonin (Biosynth, D-3203) in lysis buffer (20 mM Tris-HCl, pH 7.4, 0.1 mM EDTA, 50 mM NaCl, 10% glycerol) supplemented with 1 mM PMSF (Aladdin, P105539) and 1× homemade Protease Inhibitor Cocktail. Protein concentration was determined using BCA protein assay kit (Abbkine, KTD3001) before electrophoresis on 10% SDS or 4–12% blue native polyacrylamide gels. For immunoblotting, proteins were transferred onto PVDF membrane (Immobilon-P, 0.45 µm, Millipore, IPVH00010) and detected with primary and secondary antibodies (listed in Table S2). The blots were then developed using ECL reagents (Epizyme Biotech, SQ201L; Advansta, K-12045-D50) on the ChampChemi910 Chemiluminescent Imaging System (Beijing Sage Creation).

**Bathophenanthroline assay.** The bathophenanthroline assay was modified according to previous studies^14,15^. Cell lysate obtained using 1% DDM in ddH_2_O was added into bathophenanthroline buffer to reach a final concentration of 100 mM Tris-HCl (pH 7.4), 0.6% SDS and 10 mM bathophenanthroline (Macklin, B828385). To determine Fe^2+^ concentration, cell lysate was directly added into the bathophenanthroline buffer. For total iron (Fe^2+^ and Fe^3+^) measurement, Fe^3+^ in the cell lysate was reduced by 100 mM DTT at 80°C for 10 minutes. The standard curve was prepared using ammonium ferrous sulfate hexahydrate (Aladdin, A112646). To ensure that both Fe^2+^ and Fe^3+^ ions could be measured unambiguously in the same sample, ammonium ferrous sulfate was mixed with ammonium ferric citrate (Aladdin, A100170) at different ratios with a total iron concentration of 500 μM.

**Oxygen consumption rate (OCR) measurement.** Cells were seeded at a previously determined optimized density (~20,000 cells per well) in the Seahorse XF Cell Culture Microplates (Agilent, 102601-100). The OCR over time was monitored by Agilent Seahorse XF Analyzer at untreated condition followed by injection of oligomycin (1.5 µM), FCCP (1 µM for PaKiT, 0.5 µM for HEK293T) and rotenone/antimycin A (0.5 µM) from Seahorse XF Cell Mito Stress Test Kit (Agilent, 103015-100). Subsequently, cells were stained by 5 μg/mL Hoechst 33342 (Sigma, B2261) for 10 min at 37 °C to determine cell density for normalization. Results were analyzed using Agilent Seahorse XFe96 and XFe24 Analyzers Wave 2.6 and GraphPad Prism 6 softwares.

**RNAseq and GO enrichment analysis.** PaKiT cells were treated in triplicate with 50 µM rotenone for 6 h or 20 μM VBIT-4 for 24 h using DMSO as control before lysed in TRIzol reagent (Vazyme, R401-01) on ice. Lysates were transferred to RNase- and DNase-free microcentrifuge tubes to extract RNA. 200 µL chloroform (SCR, 10006818) was added to 1 mL TRIzol lysate before mixing thoroughly. Samples were incubated for 10 min on ice before centrifugation by 12,000 ×g for 15 min at 4 °C. Samples were separated into three layers and the transparent upper layer was transferred into another RNase- and DNase-free microcentrifuge tube before adding equal volume of ice-cold isopropyl alcohol (SCR, 80109218) and mixing gently by inverting. Samples were incubated for 10 min on ice before centrifugation by 12,000 ×g for 10 min at 4 °C. The RNA pellet was washed in 75% ethanol (SCR, 10009218) prepared with diethyl pyrocarbonate (DEPC)-treated water (Sigma, D5758) and finally dissolved in DEPC-treated water. RNA quality was examined by gel electrophoresis with Qubit (Thermo). Libraries were constructed using VAHTS Stranded mRNA-seq Library Prep Kit for Illumina (Vazyme, NR602), and sequencing was carried out using the Illumina Novaseq 6000 instrument by the commercial service of Genergy Biotechnology Co. Ltd. (Shanghai, China). The raw data was handled by Skewer and data quality was checked by FastQC (v0.11.2). Clean reads were aligned to the reference genome using STAR (2.5.3a). Afterwards, a differential gene expression analysis was performed using DESeq2 R package^43^. For GO enrichment analysis, mitochondrial related genes were defined according to MitoCarta 3.0 and MitoCoP databases^44,45^. Differentially expressed genes were subjected to Metascape analysis^46^.

**Metabolomic profiling** PaKiT cells were pre-treated with 50 μM ruthenium red for 1 day before 10 μM rotenone treatment for 1 hour followed by metabolomics analysis. PaKiT cells were calculated before centrifugation, and the pellet was flash-frozen in liquid nitrogen and stored in -80 °C before metabolomic profiling. Cell pellets were lysed with lysis buffer (methanol:acetonitrile:water 5:3:2 *v/v/v*) to the concentration of 1 × 10^7^ cells/ mL. Then the suspensions were vortexed for 30 minutes and centrifuged at 18,213 g for 10 minutes at 4°C. Supernatants were injected into the Ultra-High-Pressure Liquid Chromatography–Mass Spectrometry (UHPLC-MS) system. Samples were analyzed using a 5-minute gradient as previously described^47,48^. In brief, metabolites were separated on a Kinetex C18 column (2.1×150 mm, 1.7 um, Phenomenex) by the following chromatography conditions: flow rate 450 μl/min, column temperature 45°C, and sample compartment temperature 7°C. Positive mode was from 5% to 95% of ACN/0.1% Formic Acid in Water/0.1% Formic Acid and negative mode was from 95% ACN/5% water/1mM ammonium acetate in 5% ACN/95% water/1mM ammonium acetate. Samples were randomized and run in positive and negative ion modes independently. Raw data files were converted to mzXML format using RawConverter and analyzed via Maven. The metabolomic data were normalized based on website MetaboAnalyst (<https://www.metaboanalyst.ca/MetaboAnalyst/>).

**Fatty acid preparation**

Linoleic acid was dissolved in DPBS containing 100 mg/ml BSA to prepare a stock solution with a final concentration of 3.2 mM. The control group was supplemented with an equivalent amount of DPBS with BSA.

**Lipid droplet size and intensity analysis.** Lipid droplet signal intensity and size were calculated after staining with 1 μg/mL BODIPY 493/503 (Macklin, D849506) in DMEM/F-12 media without FBS. Living cell images of each sample were acquired and analyzed using Fiji software. Briefly, the cell edges were manually outlined to measure the average fluorescent intensity of BODIPY 493/503 in individual cells. To calculate the size of lipid droplets, three largest lipid droplets in each cell were selected to measure their areas using the circular tool.

**Statistical analysis.** Statistical analysis was performed using GraphPad Prism 6. Significance between two groups was determined by two tailed Student’s *t*-test. Significance for pairwise comparison among multiple groups was determined by one-way ANOVA with Dunnett's test or Tukey's test. *, *P* < 0.05; **, *P* < 0.01; ***, *P* < 0.001; ****, *P* < 0.0001; ns, not significant. Data were shown as mean ± s.d., mean ± s.e.m. or box plot. For multiple comparisons issue, we adjusted *P*-values based on the Benjamini and Hochberg (BH) method^49^ and ensured that the false discovery rate (FDR) < 0.05.
